# Indirect decoding of behavior from brain signals via reconstruction of experimental conditions

**DOI:** 10.1101/2022.03.24.485588

**Authors:** Joram Soch, John-Dylan Haynes

## Abstract

The data acquired during a functional magnetic resonance imaging (fMRI) experiment usually comprise experimental conditions, brain signals and behavioral responses. This reflects the underlying causal flow where the experimental conditions evoke brain responses that in turn result in behavior. In multivariate analyses, a common approach is to focus on the second step of that chain and to decode behavioral responses from brain signals. However, a different approach would be to first reconstruct the experimental conditions from brain signals and in turn use these to predict behavior. While this indirect approach would go against the causal chain of events, it might work better under certain circumstances, especially when the experimental conditions evoke much stronger measurable brain signals than the overt motor behavior. Here we tested this question directly by assessing the various mappings between conditions, brain signals and behavior in an open dataset. We found that the path of first decoding experimental conditions works surprisingly well, even though in our example data set, it is still outperformed by directly decoding the behavior from brain responses.

## 1 Introduction

In cognitive neuroscience, relationships between mind and brain are frequently investigated using neuroimaging experiments, i.e. by conducting a psychological task with human participants and measuring their brain activity when performing the task (e.g. via fMRI or EEG). The information acquired and stored during such neuroscientific experiments can usually be categorized into *experimental design* X (e.g. experimental conditions, modulator variables), *physiological signals* Y (i.e. hemodynamic activity, electric potentials) and *behavioral data* Z (e.g. button presses, stimulus ratings). The physiological signals can be further distinguished into *stimulus-related signals* Y_1_ (i.e. measured during stimulus presentation) and *response-related signals* Y_2_ (i.e. measured during behavioral responses). As conditions represent the input side and responses correspond to the output side of the system, causal influences would be assumed to flow from experimental conditions to stimulus-related signals, to response-related signals, and finally, to behavioral responses (X *→* Y_1_ *→* Y_2_ *→* Z).

Conventional neuroimaging data analysis typically aims at using experimental conditions or behavioral responses to predict measured signals (“generative models”, e.g. general linear models; Friston et al., 1994; Friston and Price, 2001) or to decode experimental inputs or behavioral outputs from measured signals (“decoding algorithms”, e.g. support vector machines; Cox and Savoy, 2003; Haynes, 2015). For example, in multivariate pattern analysis (MVPA) for functional magnetic resonance imaging (fMRI), decoding algorithms can be trained to predict behavioral responses from measured signals (e.g. Soon et al., 2008) or to reconstruct experimental design variables from measured signals (e.g. Reverberi et al., 2012a, b). Similarly, general linear models (GLMs) for fMRI can model measured signals as a function of both, actual stimulus properties and/or a subject’s later responses (e.g. Soch et al., 2021).

In the case that a cognitive neuroimaging experiment involves an instructed task, with rules telling the subject how to behave, it is also possible to assess the mapping from presented stimuli to the subject’s reactions – which, in principle, allows to predict the behavioral responses from the experimental design (X *→* Z; stimulus-based response decoding, sbRD). This is what is frequently done in purely behavioral studies in experimental psychology. On the other hand, behavioral responses can also be directly decoded from fMRI signals measured during those responses (Y_2_ *→* Z; direct response decoding, dRD). In this paper, we address the question whether it is possible to decode behavioral responses from fMRI indirectly, i.e. by learning experimental conditions from fMRI signals measured during stimulus presentation and then predicting responses using a purely behavioral model (Y_1_ *→* X *→* Z; indirect response decoding, iRD).^1^

In contrast to dRD, which directly extracts behavioral responses Z from response-related fMRI signals Y_2_, iRD operates indirectly by first decoding the experimental design X from stimulus-related fMRI signals Y_1_ and then predicting behavioral responses from the experimental design using the mapping from X to Z. *In this way, behavioral responses are decoded from fMRI signals without learning them directly from those signals*.

At first sight, one might assume that direct response decoding should in principle work better than indirect response decoding, given that a mapping Y *→* X *→* Z (iRD) is a special case of the mapping Y *→* Z (dRD) and that brain activity Y is the proximal cause of a subject’s behaviour, whereas experimental conditions X are only a distal cause. However, there are reasons why this might not be the case. First, brain signals Y_1_ predictive of X might be different from brain signals Y_2_ predictive of Z, i.e. certain conditions might favor prediction of Z from Y_1_ *→* X rather than directly from Y_2_. Second, the mapping Y_1_ *→* X extracts the behaviorally relevant features (i.e. experimentally controlled variables such as experimental conditions and stimulus properties) from fMRI signals and there should be a direct mapping X *→* Z (e.g. Görgen et al., 2018), if subjects are instructed to react with specific responses to specific conditions (e.g. Hebart et al., 2012). Thus, the question which approach is more precise is not as simple as it may seem.

In order to answer this question, we analyzed a large fMRI data set (Botvinik-Nezer et al., 2019) using these different strategies. We chose this particular data set, because it allows to conceptualize experimental variables X and behavioral responses Z in a number of different ways (see Section 3.4), thereby allowing to assess robustness of results. All in all, we find that, as one might expect, dRD mildly outperforms iRD, but both allow for above-chance decoding of behavioral responses (see Section 4.1).

The present paper is organized as follows: First, we introduce the theoretical framework underlying dRD and iRD by defining the respective mappings between X, Y_1_/Y_2_ and Z (see Section 2). Second, we describe a large empirical data set and the behavioral and neurophysiological models applied to those data (see Section 3). Then, we report decoding accuracies of dRD and iRD and how analysis choices influence the performance of iRD (see Section 4), before we finally discuss our results (see Section 5).

Along with this paper, we also provide supplementary material inlcuding an empirical pilot study using a smaller data set (see Supplementary Section 1) and extended results using the large data set (see Supplementary Section 2).

## 2 Theory

In this section, we introduce the theoretical framework behind direct and indirect response decoding. Non-technical readers are recommended to have a look at the graphical summary of the methodology (see Figure 1) and then directly proceed to the empirical data analysis (see Section 3).

**Figure 1:**
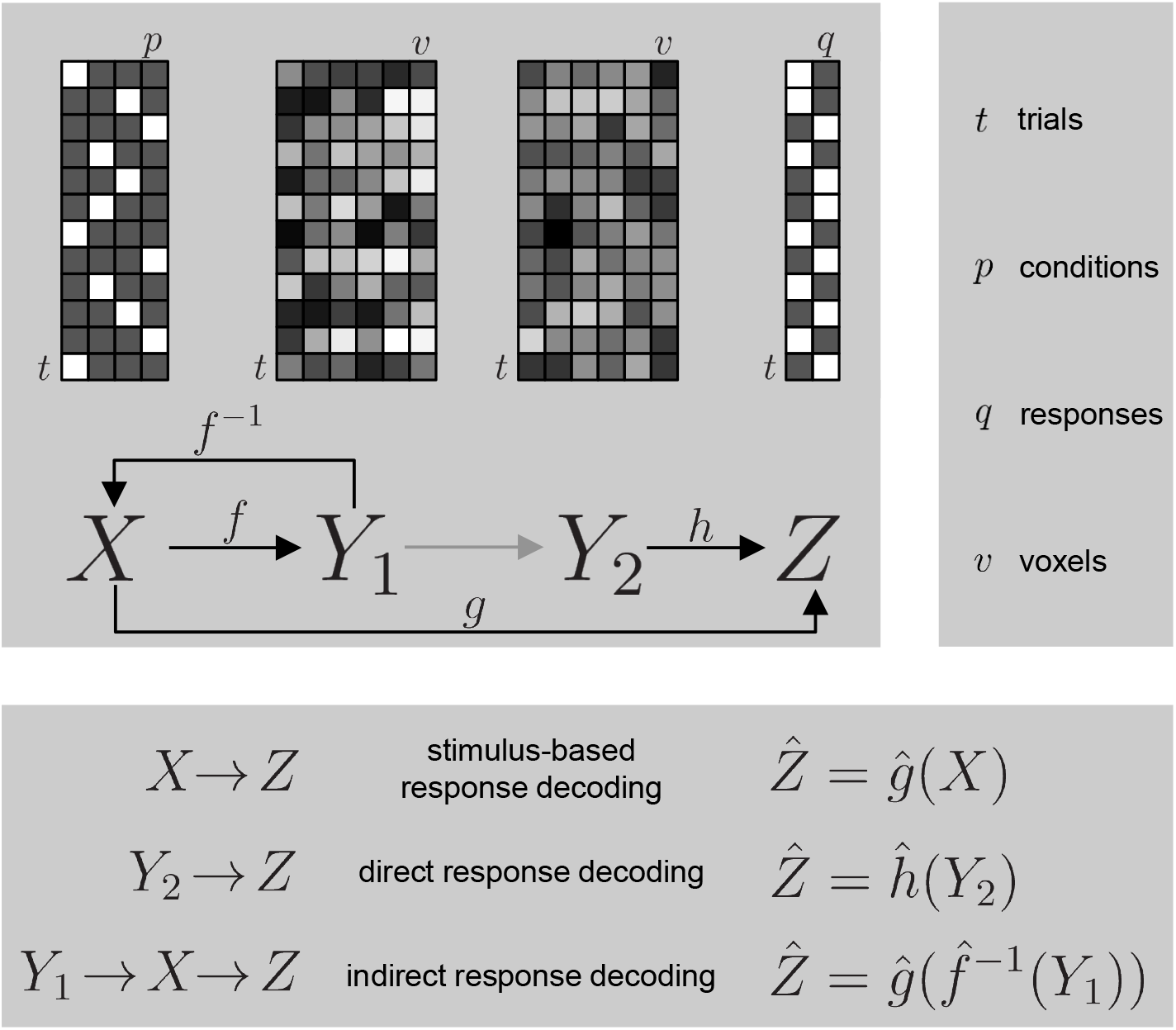
Theory behind indirect response decoding. The analyses operate on the trial-bycondition matrix *X* (experimental design), the trial-by-voxel matrices *Y*_1_ and *Y*_2_ (stimulus-related and response replated fMRI signals in e.g. searchlight, region of interest or whole brain), and the trial-by-response matrix *Z* (behavioral responses). The theory assumes a neurophysiological model *f* (from *X* to *Y*_1_, inverse *f* ^*−*1^), a psychobehavioral model *g* (from *X* to *Z*) as well as a mapping for direct response decoding *h* (from *Y*_2_ to *Z*).

### 2.1 The neurophysiological model

Consider *brain responses Y* acquired from a number of measurement channels (e.g. fMRI voxels or EEG electrodes) during a number of experimental measurements (e.g. scans or trials). Further assume *experimental variables X* that can be temporally assigned (i.e. scan-wise or trial-wise) to the measured data *Y*. For simplicity, we will assume that *Y* and *X* have already been reduced to the level of trials, e.g. by estimating trial-by-trial response amplitudes using trial-wise hemodynamic response functions (HRF; Mumford et al., 2012) or by analyzing time-locked event-related potentials (ERP; Roach and Mathalon, 2008), such that *Y* is a *t × v* matrix and *X* is a *t × p* matrix where *t* is the number of trials, *v* is the number of channels and *p* is the number of predictors.

Then, a *neurophysiological (forward) model* is defined as a mapping *f* from the experimental design matrix to the measured data matrix (cf. Friston et al., 2008, eq. 8)

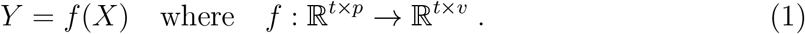

For example, if trial-wise parameter estimates and trial-by-trial covariance matrix were obtained via inverse transformed encoding models (ITEM; Soch et al., 2020), the neurophysiological model would be a multivariate general linear model (MGLM) given by^2^

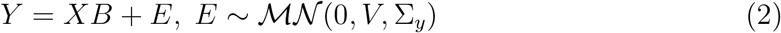

where *B* is a *p × v* matrix of voxel-wise parameters, *E* is a *t × v* noise matrix of matrix-normally distributed errors, *V* is the known temporal (trial-by-trial) covariance matrix and Σ_*y*_ is the unknown spatial (voxel-by-voxel) covariance matrix. If the goal was to infer on the parameters *B*, the model would be estimated, e.g. via maximum likelihood (ML), and statistical tests would be performed (Allefeld and Haynes, 2014).

Given the forward model in (1), we can also conceive an *inverse neurophysiological model*, mapping from measured data to experimental design (cf. Friston et al., 2008, eq. 10)

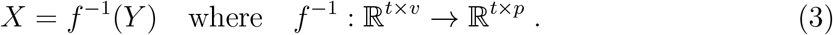

For example, for the multivariate GLM given in (2), the inverse model is also an multivariate GLM (Soch et al., 2020, eqs. 10 and C.2) given by

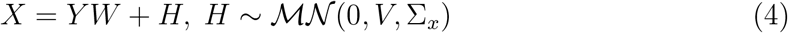

where *W* is a *v × p* weight matrix, *H* is a *t × p* error matrix and Σ_*x*_ is the condition-by-condition covariance matrix which is given by Σ_*x*_ = *W* ^T^Σ_*y*_*W*.

Note that so far, we are quite generally refering to “brain signals” *Y*. Although the neurophysiological model is primarily designed for modeling stimulus-related signals *Y*_1_ in terms of the experimental design *X* (or predicting *X* from *Y*_1_), the formalism given by (1) and (3) can in principle also be used for modelling response-related signals *Y*_2_ in terms of behavioral responses *Z* (or predicting *Z* from *Y*_2_; see Section 2.4).

In Table 1, we list several forward and inverse neurophysiological models for illustration.

**Table 1:** Examples of neurophysiological models. Various forward and inverse models are listed with their core equations and exemplary applications in neuroimaging. Abbreviations: GLM = general linear model, MGLM = multivariate GLM, SVC = support vector classification, SVR = support vector regression.

| Category | Name | Equation | Reference |
| --- | --- | --- | --- |
| forward | univariate GLM | $y = X\beta + \varepsilon, \varepsilon \sim \mathcal{N}(0, \sigma^2 V)$ | Friston et al., 1994 |
| forward | multivariate GLM | $Y = XB + E, E \sim \mathcal{MN}(0, V, \Sigma_y)$ | Allefeld and Haynes, 2014 |
| inverse | inverse MGLM | $X = YW + H, H \sim \mathcal{MN}(0, V, \Sigma_x)$ | Soch et al., 2020 |
| inverse | logistic regression | $\log \frac{p_i}{1-p_i} = y_i w + \varepsilon_i, p_i = \Pr(x_i = 1)$ | Mumford et al., 2012 |
| inverse | linear SVC | $x_i = \text{sgn} [wy_i^T + b], x_i \in \{-1, +1\}$ | Cox and Savoy, 2003 |
| inverse | linear SVR | $x_i = [wy_i^T + b], x_i \in \mathbb{R}$ | Heinzle et al., 2011 |

### 2.2 The psychobehavioral model

Assume that, in parallel to the brain signals *Y*, we have also measured *behavioral responses Z* represented as a *t × q* matrix where *q* is the number of recorded behavioral variables, e.g. button presses or stimulus ratings. Unless the subject’s behavior is intrinsically driven (Soon et al., 2008), the behavioral responses *Z* should, just like the brain signals *Y*, in some way also depend on the experimental design variables *X*.

Thus, a *psychobehavioral (forward) model* (PBM) is defined as a mapping *g* from the experimental design matrix to the behavioral data matrix:

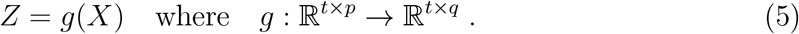

Such a PBM can be arbitrarily complex, ranging from linear regression capturing mean differences in reaction times (*Z*) between experimental conditions (*X*) to a Bayesian hierarchical model describing learning under uncertainty (Mathys, 2011).

For illustrative purposes, let us assume that the psychobehavioral model in (5), similar to the neurophysiological model in (2), is also a general linear model, but with independent and identically distributed (i.i.d.) trials (*V* = *I*_*t*_), i.e.

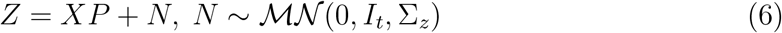

where *P* is a *p × q* mapping matrix, *N* is a *t × q* noise matrix, *I*_*t*_ is the *t × t* identity matrix and Σ_*z*_ is the response-by-response covariance matrix.

For example, if *Z* and *X* are both continuous (e.g. reaction times and stimulus intensities), the coefficients in *P* represent linear effects from *X* on *Z* (and *X* may include a constant regressor to allow for non-zero intercepts); if *X* becomes a binary indicator variable (e.g. experimental conditions), the entry *p*_*ij*_ represents the mean of response variable *j* within experimental category *i*; and if *Z* also becomes an indicator variable (e.g. button presses), the entry *p*_*ij*_ represents the transition probability Pr(*z*_*j*_ = 1|*x*_*i*_ = 1) of chosing the response category *j*, given experimental category *i*.

In Table 2, we list examples for those different types of psychobehavioral models.

**Table 2:** Examples of psychobehavioral models. For each combination of continuous or discrete behavioral variables and design variables, a possible analysis method is given.

| Type | Behavior $Z$ | Design $X$ | Analysis |
| --- | --- | --- | --- |
| continuous–continuous | reaction times | stimulus intensities | linear regression |
| continuous–discrete | stimulus ratings | stimulus types | analysis of variance |
| discrete–continuous | option accepted | motivational value | logistic regression |
| discrete–discrete | item remembered | emotional valence | conditional probabilities |

### 2.3 Stimulus-based response decoding

Given such a psychobehavioral model, *stimulus-based response decoding* usually consists in estimating the model – most often a linear model such as (6) –, and testing model parameters for statistical significance.

If the focus is on prediction rather than inference, the analysis would proceed by estimating the function *g* from training data

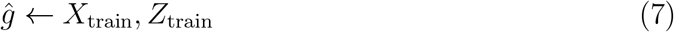

and applying it to test data

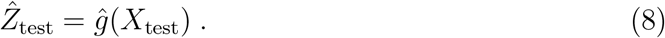

Here, we perform leave-one-run-out cross-validation (CV) over fMRI recording sessions which allows to derive behavioral predictions for the entire data *Z*. More precisely, in the *i*-th CV fold, the session *i* serves as the test set and sessions *j* = 1, …, *S, j* ≠ *i* serve as the training set, where *S* is the number of sessions.

This procedure can also be referred to as “*decoding behavior from the design*” (e.g. Görgen et al., 2018, Fig. 4), because the behavioral responses *Z* are predicted solely from the experimental design *X*, without recourse to the measured signals *Y*.

### 2.4 Direct response decoding

If the signals *Y*_2_ measured during the subject’s responses are to be taken into account, then behavioral responses *Z* – whether continuous or discrete (see Table 2) – can simply be treated similar to the experimental variables *X* and the goal is to find a function *h* mapping from *Y*_2_ to *Z*:

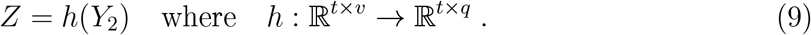

This is analogolous to an inverse neurophysiological model such as (3) or similar (see Table 1), but with behavior *Z* instead of design *X* as the target variable and with response-related signals *Y*_2_ as sourve variables.

Again, *h* would be estimated from training data

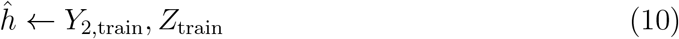

and applied for prediction in the test data

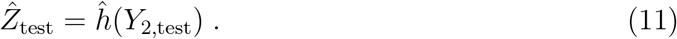

Once more, cross-validated application of (10) and (11) yields behavioral predictions for the entire data *Z* based on independent parts of *Y*_2_.

We refer to this procedure as “*direct response decoding*” (dRD), because it represents the most straightforward and most common approach of predicting behavioral responses *Z* from measured signals *Y* in fMRI decoding.

In Table 3, we list examples for such direct response decoding analyses.

**Table 3:**
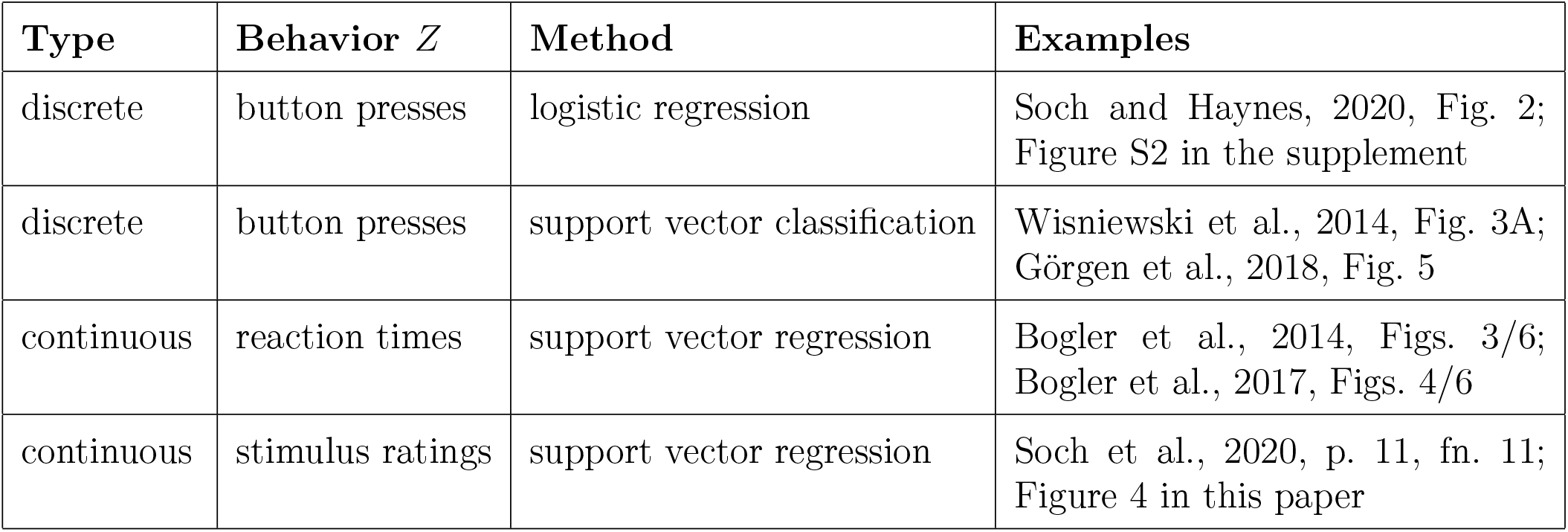
Examples for direct response decoding. For several types of behavioral response variables, a possible analysis method and examples are given.

### 2.5 Indirect response decoding

Instead of directly retrieving the behavioral responses *Z* from the experimental design *X* (see Section 2.3) or from response-related signals *Y*_2_ (see Section 2.4), one can also take an indirect path: First, we are decoding *X* from stimulus-related signals *Y*_1_ and then, we are predicting *Z* from the reconstructed design.

To recapitulate, the inverse neurophysiological model (3) defines a mapping from *Y* to *X* (see Section 2.1) and the psychobehavioral model (5) defines a mapping from *X* to *Z* (see Section 2.2). Obviously, these two functions can be composed to yield a new function mapping from *Y*_1_ to *Z* (see Figure 1):

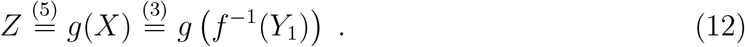

Consequently, we can estimate *f* ^*−*1^ and *g* from training data

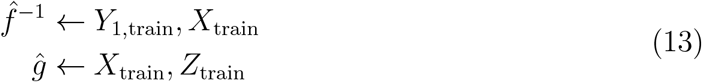

and transfer the estimated functions to test data

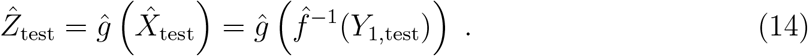

Once again, by cross-validating over fMRI runs, out-of-sample parts of *Z* are *decoded* from within-sample parts of *X* and *Y* in each cross-validation fold, yielding predictions for the entire behavioral data *Z* (see Figure 2).

**Figure 2:**
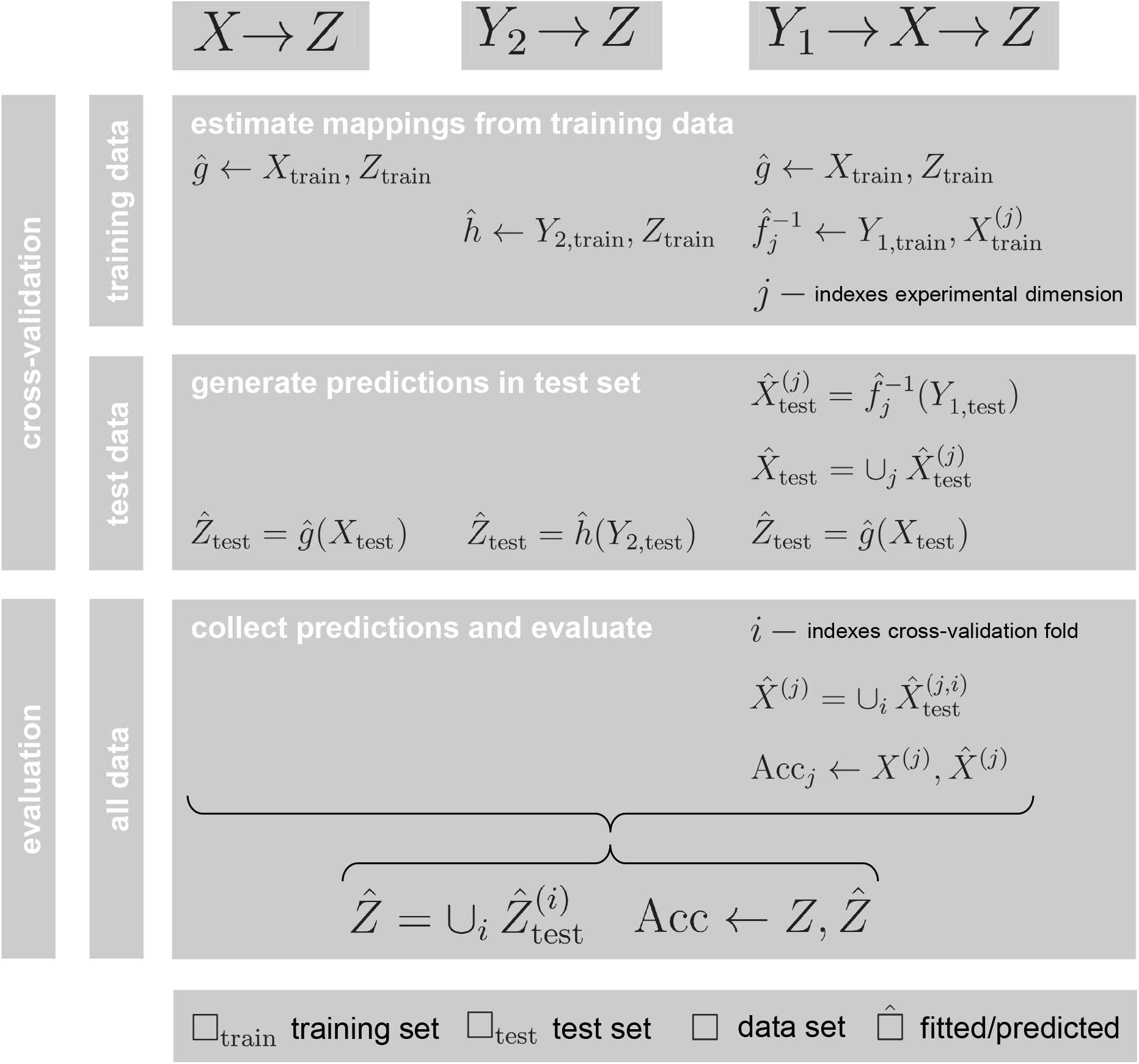
Performance assessment for response decoding. Behavioral responses are predicted separately for each cross-validation (CV) fold. For stimulus-based response decoding (sbRD, left) and direct response decoding (dRD, middle), the mapping is estimated from the training data and used to generate predictions on the test data. In indirect response decoding (iRD, right), the reconstructed design matrix 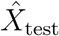 is established by combining reconstructed experimental factors (indexed by *j*) and then used for predicting behavioral responses. Finally, decoding accuracy (Acc) is assessed by collecting predicted responses 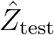 across CV folds (indexed by *i*) and comparing them to actual behavioral responses *Z*. For iRD, decoding accuracy is also calculated for predicted design variables 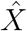, by comparing them with actual experimental variables *X*.

Whereas direct response decoding uses a mapping *Y*_2_ *→ Z*, indirect response decoding works along the path *Y*_1_ *→ X → Z*. Using iRD, button presses or stimulus ratings are not predicted directly from the measured fMRI signals, but on an indirect route through the experimental design variables (see Figure 1).

### 2.6 Experimental design dimensions

A systematic difference between dRD and iRD is that dRD directly returns predictions of the behavioral variables *Z* whereas iRD first results in predictions of the design variables *X* – which has consequences for how the decoding proceeds. To understand this, consider the following example: Assume a two-factorial design (A vs. B and 1 vs. 2) with two response options (e.g. “left” and “right”), such that *X* is a *t ×* 4 indicator matrix coding the experimental conditions A1, A2, B1 and B2; and *Z* is a *t ×* 2 indicator matrix coding the button presses C1 and C2. Then, we find that

- dRD directly predicts the behavioral dimension 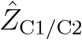 in each CV fold which can then be compared to the actual responses to calculate a measure of decoding performance;
- iRD, when experimental design dimensions are decoded separately, generates one set of predictions 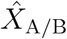 and one set of predictions 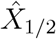; thus, these decodings have to be combined into a full design matrix^3^ in order to provide an input to the behavioral model *Z* = *g*(*X*) which then goes to predict the behavioral dimension 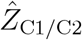.

Despite these differences, decoding performance is assessed in the same way for both approaches (and sbRD): out-of-sample predictions 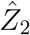 from all CV folds are collected into one set of predicted responses 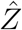 which is compared to the actual responses *Z* (see Figure 2) using a measure of decoding accuracy (DA; for discrete behavioral responses) or decoding precision (DP; for continuous behavioral responses).

## 3 Methods

Before the analysis of a large empirical data set (this section), we tested the methodological framework described here using a smaller data set. Results from this pilot analysis are reported in the supplementary material (see Supplementary Section 1).

### 3.1 NARPS fMRI data set

For the main part of this study, we used an openly available fMRI data set (*N* = 108 subjects) from the Neuroimaging Analysis Replication and Prediction Study^4^ (NARPS; Botvinik-Nezer et al., 2019, 2020).

During this experiment, subjects were offered a mixed gamble in each trial, to either win a certain amount of money or to loose a certain amount of money, with equal probability (see Figure 3). Subjects then indicated favorability of the bet using a four-point Likert scale (Likert, 1932) ranging from “strongly reject” (1) over “weakly reject” (2) and “weakly accept” (3) to “strongly accept” (4).

**Figure 3:**
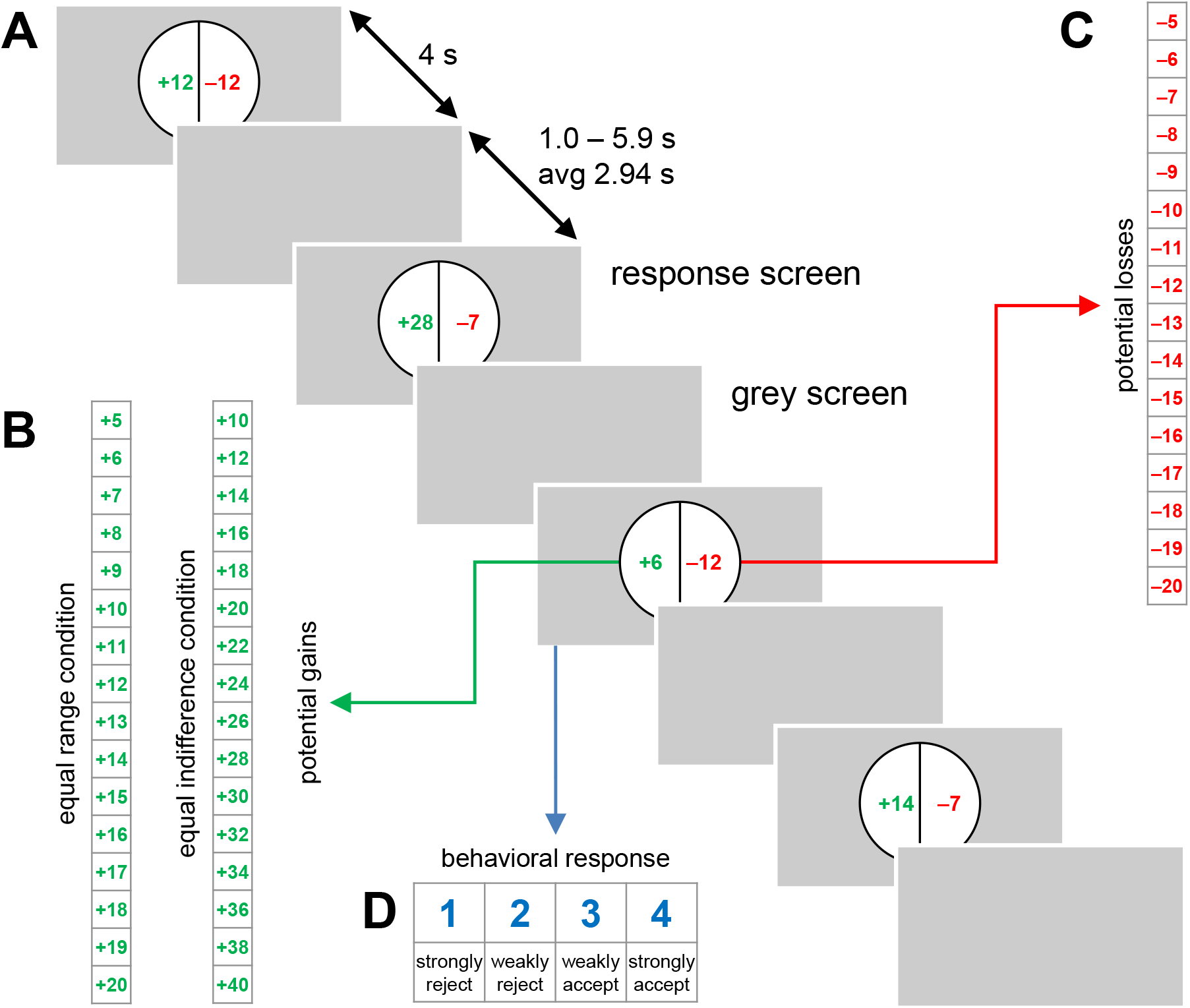
Experimental design of mixed gambling task. **(A)** In each trial, a certain amount of money to win (green) and a certain amount of money to loose (red) were displayed. Subjects had four seconds to respond whether to accept the gamble. After that, a variable inter-trial interval followed. **(B)** Potential gains in the equal range (left) and equal indifference (right) condition. Experimental condition was a between-subject factor. **(C)** Potential losses, in both experimental conditions. Each combination of potential gain and potential loss was presented exactly once to each subject, resulting in 16 *×* 16 = 256 trials. **(D)** In each trial, subjects made their response on a four-point scale.

Consequently, the experiment provides experimental design variables *X* (gain and loss), behavioral response variables *Z* (favorability) and measured fMRI signals *Y* (voxel-wise trial-wise fMRI responses). For more details, see the data descriptor (Botvinik-Nezer et al., 2019) and the description of behavioral models (see Section 3.4).

We chose this particular data set for comparing iRD vs. dRD, because (i) behavioral responses in the involved cognitive task are neither deterministic nor random with respect to the experimental design and because (ii) experimental variables as well as behavioral data can be employed as discrete as well as continuous variables (see Table 4), thereby allowing to assess the robustness of our finding.

**Table 4:** Analyses used for indirect response decoding. Both, experimental design variables and subjects’ behavioral responses, can be regarded as categorical or parametric, giving rise to four different analyses using different behavioral models.

| Analysis | Design $X$ | Signals $Y$ | Behavior $Z$ | Behavioral model |
| --- | --- | --- | --- | --- |
| 1 | high/low gain<br>high/low loss | whole-brain<br>fMRI signals | accept/reject | conditional probabilities |
| 2 | gain value<br>loss value | whole-brain<br>fMRI signals | accept/reject | logistic regression |
| 3 | high/low gain<br>high/low loss | whole-brain<br>fMRI signals | favorability rating | linear regression |
| 4 | gain value<br>loss value | whole-brain<br>fMRI signals | favorability rating | linear regression |

### 3.2 Preprocessing of fMRI data

For the analyses in this paper, we used the preprocessed images supplied with the data set on OpenNeuro^5^, generated using fMRIprep, the robust preprocessing pipeline for functional MRI (Esteban et al., 2019). For more details, see the fMRIprep protocol file available in the corresponding repository^6^.

After preprocessing, GZIP files were unpacked and normalized into a voxel size of 3 *×* 3 *×* 3mm using SPM12. Finally, a gray matter (GM) mask was generated for each subject by inclusively masking the normalized fMRIprep brain mask with SPM’s tissue probability map (TPM) for GM, thresholded at *p >* 1*/*3.

### 3.3 Statistical analysis of fMRI data

Two voxel-wise general linear models (GLMs) were fit to the fMRI data measured during each of the *S* = 4 sessions acquired from each of the *N* = 108 subjects. In one GLM, the stimulus phase of each trial was modelled using a canonical hemodynamic response function (cHRF), placed at the trial onset. This model was later used to estimate stimulus-related signals *Y*_1_ for decoding the experimental design (iRD). In the other GLM, the response phase of each trial was modelled using a cHRF placed at the time of the subject’s button press. This model was later used to estimate stimulus-related signals *Y*_2_ for decoding behavioral responses (dRD). In each model, the respective other trial phase was included as a nuisance condition.

After these models were specified in SPM12, the ITEM toolbox^7^ (Soch et al., 2020) was used to estimate voxel-and-trial-wise response amplitudes *Y*_1_/*Y*_2_ serving as feature set in the inverse neurophysiological model (see eq. 3). Finally, *The Decoding Toolbox*^8^ (Hebart et al., 2015) was used (i) to decode the experimental design using support vector machines (SVM) trained on whole-brain fMRI signals *Y*_1_ estimated using the stimulus model (for iRD) and (ii) to decode behavioral responses using SVMs trained on whole-brain fMRI signals *Y*_2_ estimated using the response model (for dRD).

### 3.4 Predicting behavioral responses from the design

When it comes to decoding behavior directly from the design, two choices can be made in the context of the present experiment. The first is whether to predict from parametric regressors (raw gain and loss, amount of money to win or loose) or to categorize those variables (high vs. low gain, high vs. low loss). The second is whether to predict parametric responses (Likert-scale responses) or to categorize those variables (accept vs. reject). In other words, we can opt for “parametric” or “categorical” at both ends of the psychobehavioral model (see Table 2).

Here, we decided to explore all these four options in order to see how stimulus-based and indirect response decoding compare to direct response decoding, depending on the form in which experimental design information is supplied to them. When design and behavioral variables are both categorical, the mapping between them can be described by simple transition probablities; when the design variables are parametric, logistic regression can predict the accept/reject decision from continuous gain and loss; finally, when the behavioral response is parametric, linear regression was used to predict the favorability rating from discrete or continuous gain and loss (see Table 4).

### 3.5 Decoding behavioral responses from fMRI signals

Direct response decoding (dRD) proceeds by decoding behavioral responses from fMRI signals. Here, we use whole-brain linear support vector machines (SVM) with a cost parameter fixed at *C* = 1, as implemented in *The Decoding Toolbox* (TDT; Hebart et al., 2015), to predict categorical and parametric responses.

In Analyses 1 and 2 (see Table 4), the categorical accept/reject dimension (response 1/2 *→* reject; response 3/4 *→* accept) is decoded from response-related fMRI amplitudes using support vector classification (SVC) and performance is quantified via decoding accuracy. In Analyses 3 and 4 (see Table 4), the parametric favorability rating (responses 1-4) is decoded from response-related fMRI amplitudes using support vector regression (SVR) and performance is quantified via correlation coefficient.

### 3.6 Decoding experimental design from fMRI signals

Just like behavioral response variables can be retrieved from measured data, experimental design variables can be decoded from fMRI signals. Again, we apply whole-brain support vector machines to achieve this goal.

In Analyses 1 and 3 (see Table 4), the categorical high/low gain and loss (below mean *→* low; above mean *→* high) are decoded from stimulus-related fMRI amplitudes using linear SVC and performance is quantified via decoding accuracies. In Analyses 2 and 4 (see Table 4), the parametric gain and loss values (mean-centered amounts of money) are decoded from stimulus-related fMRI amplitudes using linear SVR and performance is quantified via correlation coefficients.

### 3.7 Decoding behavior from fMRI, via the design

Indirect response decoding (iRD) proceeds by combining (i) an inverse neurophysiological model retrieving the experimental design from measured fMRI signals (*Y*_1_ *→ X*; see Section 3.6) with (ii) a forward psychobehavioral model mapping from the experimental design to observed behavioral responses (*X → Z*; see Section 3.4), by simply composing these two mappings (see eqs. 12 and 14).

In Analysis 1 (see Table 4), high/low gain and high/low loss are decoded from trial-wise parameter estimates, the results are used to deduce the experimental condition (e.g. high gain, low loss) and the probability of accept or reject is calculated using conditional probabilities of a particular response, given a particular condition. In Analysis 2, the gain and loss values themselves are decoded and plugged into a logistic regression model describing accept/reject as a function of gain and loss during each trial. In Analysis 3, high/low gain and high/low loss are again decoded from trial-wise response amplitudes and the parametric favorability rating is calculated using a linear mapping from experimental conditions to favorability responses. In Analysis 4, the gain and loss values are again decoded and plugged into a linear regression model describing favorability ratings in terms of gain and loss amounts.

### 3.8 Performance assessment of response decoding

As explained in the theory part, caution has to be exercised when comparing performance of sbRD, dRD and iRD (see Section 2.6). Here, we used whole-brain decoding for dRD and iRD, such that all decoding approaches result in exactly one set of predictions for *Z* per cross-validation fold. More precisely:

- The performance of decoding categorical response variables (e.g. accept/reject) or design variables (e.g. high/low gain) is quantified via decoding accuracy (i.e. number of correct predictions, divided by the total number of trials) and the balanced decoding accuracy (Brodersen et al., 2010; see Supplementary Section 2).
- The performance of decoding parametric response variables (e.g. favorability rating) or design variables (e.g. gain amount) is quantified via correlation coefficient (i.e. Pearson correlation between actual and predicted values).
- For iRD, two decoded experimental dimensions are combined by decoding them separately and selecting the (joint) experimental condition which is compatible with both marginal conditions. Then, an out-of-sample reconstructed design matrix is generated and the corresponding behavioral model is applied.

## 4 Results

Decoding performance can be assessed in several ways: First, the final accuracies of the stimulus-based response decoding (sbRD), direct response decoding (dRD) and indirect response decoding (iRD) can be compared (see Section 4.1). Second, it can be investigated how the accuracy of iRD changes when some experimental design variables are not decoded, but supplied to the behavioral model (see Section 4.2). Third, it can be investigated how well the experimental design variables themselves can be decoded (see Section 4.3).

### 4.1 Decoding accuracy of direct vs. indirect response decoding

When following the procedure described above (see Section 2.6 and 3.8), the decoding analyses result in one accuracy value (for categorical response variables) or one precision value (for parametric response variables) for each decoding algorithm (sbRD, dRD, iRD) per subject. In Figure 4, we graph these values as box plots to visualize the distribution of performances as a function of experimental group, decoding algorithm and analysis type (see Table 4).

**Figure 4:**
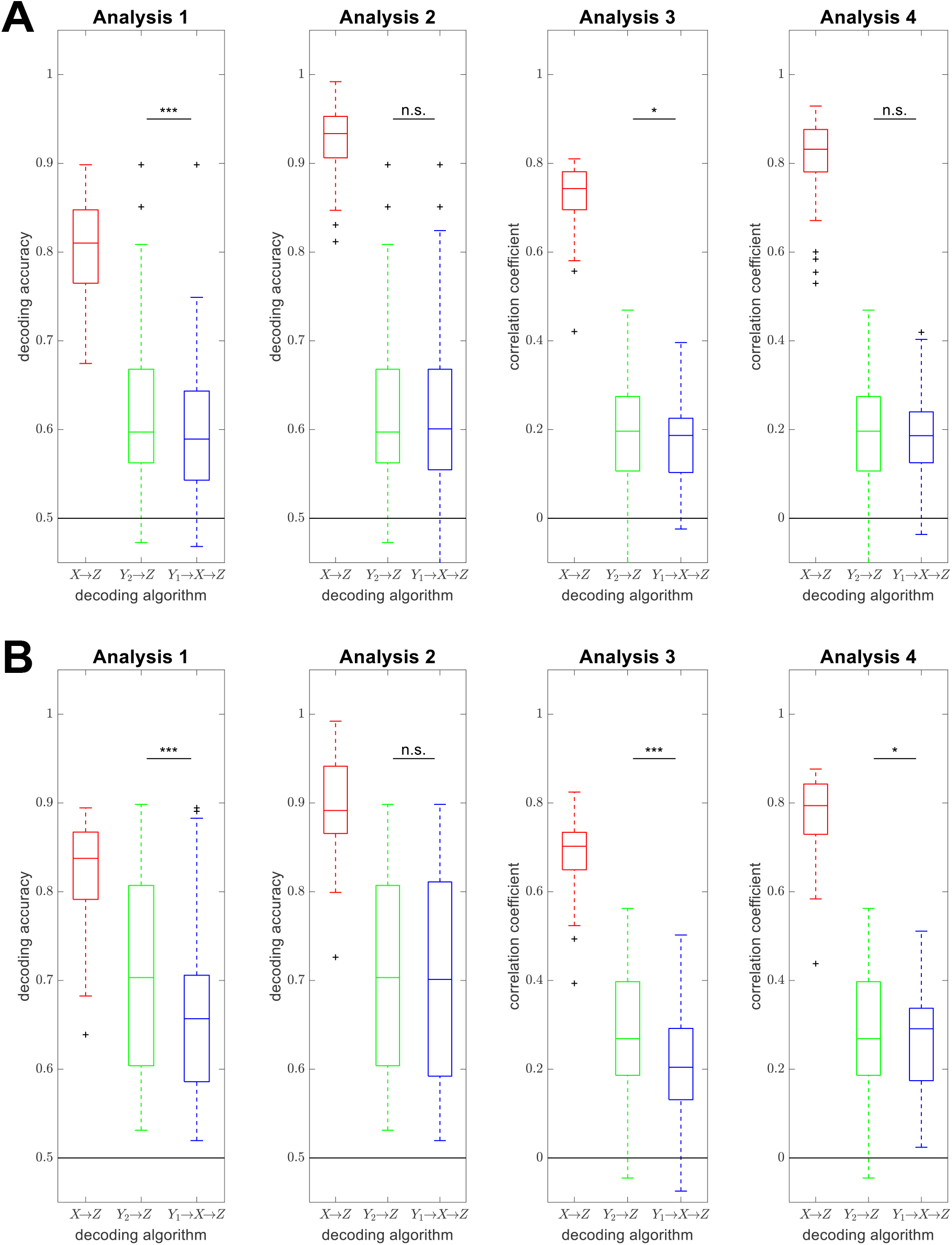
Decoding accuracies as a function of decoding method. Performances of stimulus-based (*X → Z*; red), direct (*Y*_2_ *→ Z*; green) and indirect (*Y*_1_ *→ X → Z*; blue) response decoding are visualized using box plots for four analysis types (see Table 4), separately for the **(A)** equal range and the **(B)** equal indifference condition (see Figure 3B). Direct and indirect response decoding are tested against each other using a two-tailed paired t-test. Abbreviations: n.s. = not significant; * *p <* 0.05, ** *p <* 0.01, *** *p <* 0.001.

Overall, we observe that (i) sbRD clearly outperforms dRD and iRD, because it has direct access to the experimental design and there is usually a relatively clear mapping from experimental design to behavioral responses for each subject (Botvinik-Nezer et al., 2019, Fig. 2 and 3); (ii) dRD and iRD both achieve above-chance decoding performance (accuracy larger than 0.5 and correlation larger than 0); and (iii) dRD achieves slightly higher decoding performance than iRD (see Figure 4). These observations are invariant across analysis type.

### 4.2 Decoding accuracy as a function of design information supplied

The performance of indirect response decoding can be investigated more closely by exploring how the decoding accuracy or precision changes when one or more experimental design dimensions are not reconstructed from the data, but the true values are instead supplied to the behavioral model, such that the decoding algorithm has a better knowledge of the experimental design. Regarding the present experiment, one can either (i) reconstruct the gain and loss dimension from the measured data (referred to as “iRD” in the previous section), (ii) replace reconstructed gain values by actual gain values, (iii) replace reconstructed loss values by actual loss values or (iv) supply both gain and loss values to the decoding analysis (equivalent to “sbRD” in the previous section). In Figure 5, we graph the decoding accuracies of these four approaches as a function of experimental group and analysis type.

**Figure 5:**
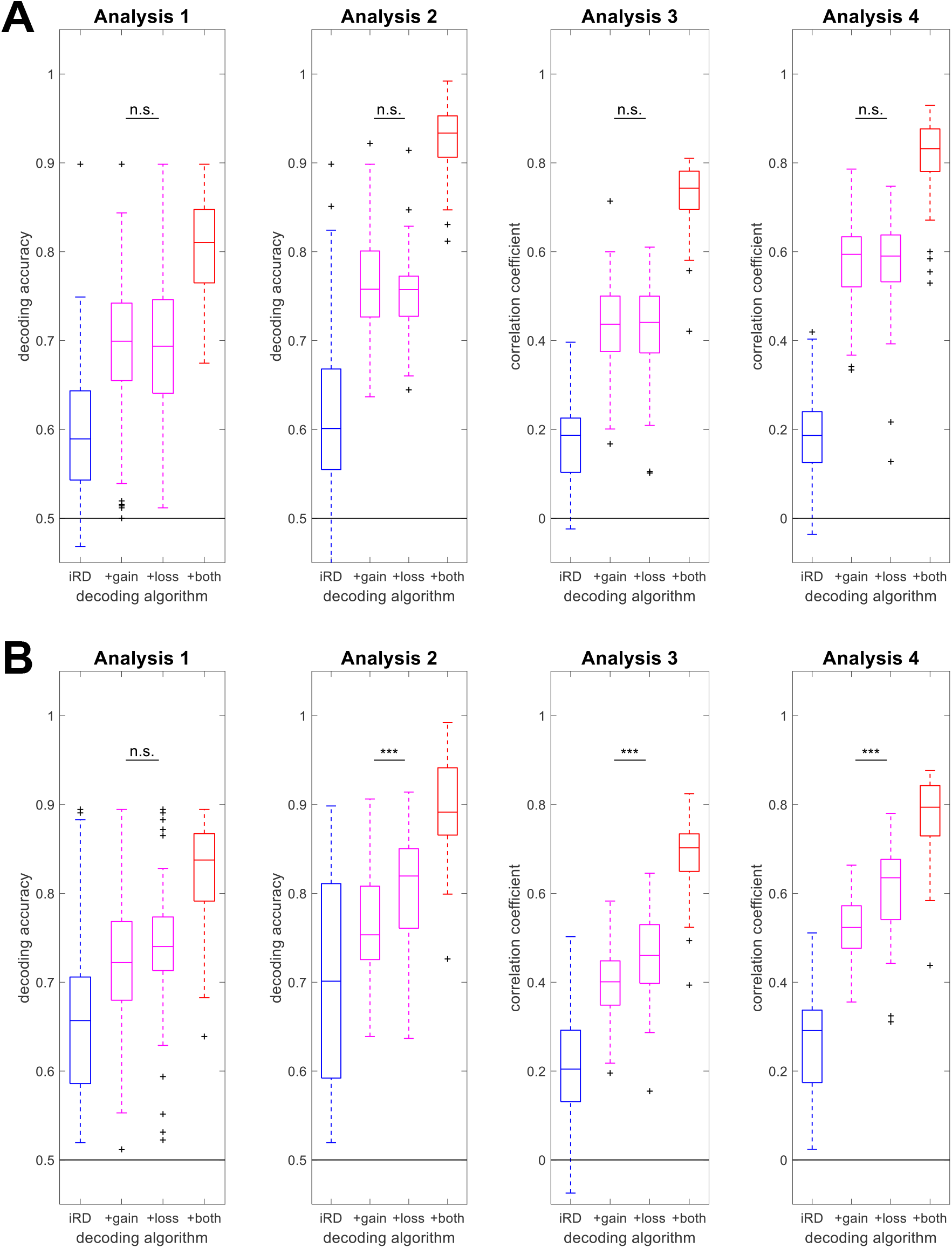
Decoding accuracies as a function of information supply. Performances of indirect response decoding (iRD), depending on whether both experimental dimensions were reconstructed from the data (blue, left), whether one of the experimental dimensions was not reconstructed, but supplied to the behavioral model (magenta, middle) or whether both experimental dimensions were known (red, right). The layout follows the one of Figure 4 and gives results for four analysis types, separately for the **(A)** equal range and the **(B)** equal indifference condition. Box plots colored in blue and red are identical to those labeled as “*Y*_1_ *→ X → Z*” and “*X → Z*” on Figure 4.

Overall, we see two different patterns: In the equal range condition, adding the true gain values and adding the true loss values increases the decoding accuracy of iRD by the same amount across analysis types (see Figure 5A), compatible with the fact that gain and loss provide the same information about subjects’ choices in this condition (Botvinik-Nezer et al., 2019, Fig. 3). In contrast, in the equal indifference condition, there was a clear linear effect, such that adding the true loss values increases the decoding accuracy of iRD more than adding the true gain values (see Figure 5B), consistent with loss being more informative than gain about subjects’ responses in this condition (Botvinik-Nezer et al., 2019, Fig. 2).

### 4.3 Decoding accuracy for experimental design variables

Finally, we can ask for the decoding accuracy when reconstructing the experimental design itself, which the indirect response decoding is based on. In the context of the present experiment, there are two experimental design dimensions (gain and loss) which either come in categorical (high/low gain/loss) or parametric (mean-centered gain/loss values) form. In Figure 6, we report decoding accuracies for experimental categories and correlation coefficients for parametric regressors as a function of experimental group and analysis type.

**Figure 6:**
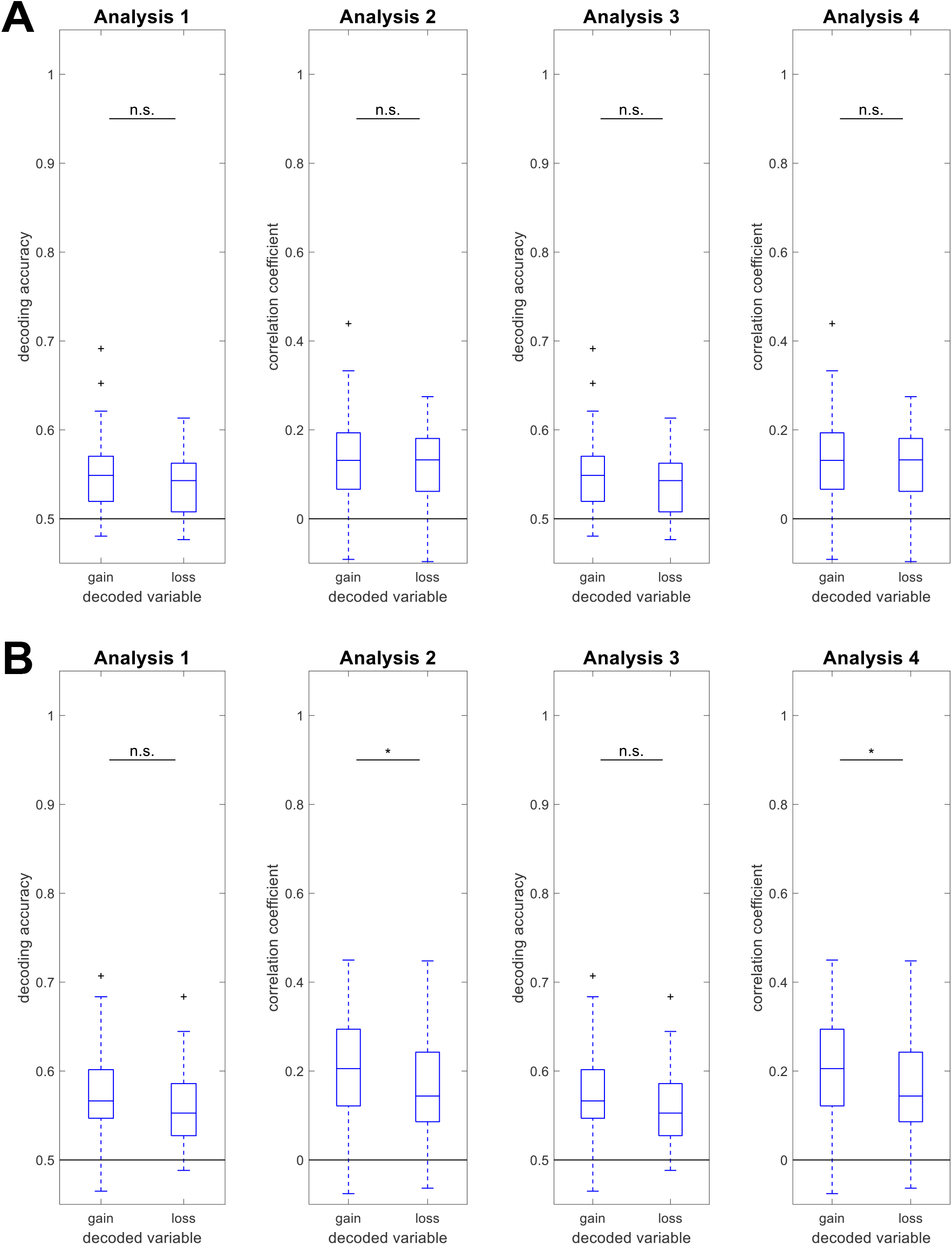
Decoding accuracies for experimental design variables. Performances of reconstructing the experimental design from measured fMRI signals. The layout follows the one of Figure 4 and gives results for four analysis types, separately for the **(A)** equal range and the **(B)** equal indifference condition.

Overall, wee see that (i) decoding accuracies for experimental design variables are relatively low (accuracies of around 0.6 and correlations of around 0.2), but (ii) gain and loss can be equally well reconstructed from trial-wise fMRI signals across analysis types in both experimental groups, equal range and equal indifference (see Figure 6). This means that, although true loss values provide more information about behavioral responses than true gain values in the equal indifference condition (see Figure 5B), measured fMRI signals provide the same amount of information about loss values and gain values in this condition (see Figure 6B).

## 5 Discussion

In this study, we have considered statistical mappings between measures for mind, brain and behavior. Specifically, we were looking at experimental design X, behavioral responses Z as well as stimulus-related fMRI signals Y_1_ and response-related fMRI signals Y_2_. We introduced several mappings between those quantities, allowing us to define *indirect response decoding* (iRD) of Z from Y_1_ via X and contrast it with *direct response decoding* (dRD) which attempts to predict Z from Y_2_ directly.

### 5.1 Performance of indirect response decoding

The results of our study can be summarized as follows:

- iRD and dRD both allow for above-chance decoding of parametric or categorical behavioral responses (see Figure 4).
- When considered across analysis strategies and experimental groups, dRD mildly outperforms iRD (see Figure 4).
- iRD levels with dRD, even when two experimental dimensions (gain and loss), but just one behavioral dimension (favorability) have to be reconstructed.
- iRD improves when experimental dimensions (gain or loss) are not decoded, but supplied to the model, depending on which dimension is supplied (see Figure 5).
- Experimental dimensions which can be decoded equally well from fMRI activity may still have different power to predict behavioral responses (see Figures 5 and 6).

Intuitively, it would not appear very promising to indirectly predict behavioral responses, because the additional data transformation step potentially introduces additional sources of noise which in turn could reduce decoding accuracy.

However, our results demonstrate that behavioral responses can be decoded without training on neurophysiological data Y_2_ measured during behavioral responses Z, but rather indirectly by taking a detour and decoding from neurophysiological data Y_1_ measured during experimental conditions X – sometimes equally well as when predicting Z directly from Y_2_ (see Figure 4). This is particularly interesting, because dRD is often seen as a “sanity check”, the decoding accuracy of which should not be reached by other analyses (see e.g. Görgen et al., 2018, Fig. 5).

### 5.2 Factors influencing indirect response decoding

This main result is stable across analysis choices for the behavioral dimension (categorical: accept vs. reject; parametric: favorability) and for the experimental dimensions (categorical: high/low gain/loss; parametric: raw gain/loss) as well as the experimental group subjects were assigned to (equal range vs. equal indifference).

It is also worth noting that in our example, just one response dimension (favorability), but two design dimensions (gain and loss) had to be decoded from fMRI (see Table 4). We therefore hypothesize that decoding the design from the data (i.e. iRD’s first step) acts as a feature reduction mechanism which helps iRD predicting behavior using the *psychologically most meaningful factors*. Potentially, this advantage over dRD cancels out with (i) the fact that iRD involves two transformations which are subject to noise (Y_1_ *→* X and X *→* Z) and (ii) subjects do not behave perfectly which reduces the accuracy of the psychobehavioral model (i.e. iRD’s second step).

Obviously, the accuracy of iRD depends on both, (i) the accuracy of the inverse neurophysiological model (Y_1_ *→* X) – the more clearly the experimental design can be reconstructed, the more psychologically meaningful information is retrieved from brain activity; and (ii) the accuracy of the psychobehavioral model (X *→* Z) – the more clearly the experimental design maps to behavioral responses, the more of the reconstructed design is carried through to behavioral predictions.

### 5.3 Thought experiment and future research

Along these lines, one can think of an example in which iRD might outperforms dRD, even if this was not observed in the present study. Because the mappings Y_1_ *→* X and Y_2_ *→* Z likely use different parts of the brain with different signal-to-noise ratios (SNR) and different discriminability of predicted variables, there might be situations in which the “stimulus SNR” is higher than the “response SNR”.

Imagine the following experiment: Subjects are stimulated with large flickering checkerboard stimuli as visual cues in either the left or the right hemifield. Their task is to press one of two buttons assigned to these two conditions, but to use one finger for both buttons. This would lead to very strong differential brain responses in early visual cortex, while the two different motor actions would nonetheless produce very similar brain responses. Additionally, because the stimuli are highly discriminable, the purely behavioral model predicting respones from stimuli would perform very well. Thus, we would observe very discriminative stimulus-related brain activity Y_1_ and very indiscriminative response-related brain activity Y_2_. Consequently, whole-brain activity would likely hold more information about conditions than about responses and the performance of iRD should be higher than that of dRD when conducting this experiment.

It should also be noted that our results circumvent the commonly assumed causal chain in which the brain (Y_1_/Y_2_) is a necessary intermediate station processing stimulus input (X) into behavioral output (Z). Thus, predicting behavior from brain activity should outperform predicting behavior from presented stimuli, as the brain is closer (more proximal) to the predicted event than the stimulus (more distal) from a physical-causal point of view. However, the reverse is usually observed (see Figure 4), most likely because the distal cause “stimulus” is measured with a much larger accuracy than the proximal cause “brain”. Whether the problem can be approached from a different perspective using a mind-brain-behavior mediation analysis, will be the subject of future research.

### 5.4 Applicability of indirect response decoding

Although this was more or less a proof-of-concept study, the presented method may be valuable in the future as some sort of *intrinsicality measurement* for a neuropsychological task. For example, when performing a perceptual decision-making task (Hebart et al., 2012; Wisniewski et al., 2014; Pischedda et al., 2017; Hebart and Görgen, 2015; Soch and Haynes, 2020; Barbieri et al., 2022), the stimulus-response mappings are in principle fixed by the instructions of the task, so that stimulus-based response decoding is expected to perform very well (at high levels of sensory evidence) and iRD is expected to predict above chance. In contrast, when subjects identify stimuli in the near-absence of visual information (at low levels of sensory evidence), when they perform an intrinsically driven task (e.g. Soon et al., 2008) or are even asked to behave unpredictably (e.g. Schultze-Kraft et al., 2016), the psychobehavioral model is not expected to explain behavior (or there might not even be experimental variables to predict from), such that iRD is expected to remain at chance level. The task used here is somewhere in between these two, as subjects were free to indicate their behavioral responses (favorability), but experimental variables (gain and loss) can be expected to have a significant influence on these responses. It remains to be seen which and how properties of the employed cognitive task influence (indirect) response decoding accuracy.

## Footnotes

1 The use of Y_1_ and Y_2_ here signifies that experimental conditions and behavioral responses are typically predicted from brain activity measured during *different phases* of the *same trial* (Y_1_: stimulus-related signals; Y_2_: response-related signals).

2 In the ITEM framework, the estimated trial-wise response amplitudes (here: *Y*) are denoted as 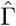 (Soch et al., 2020, eq. 10), the trial-wise experimental design (here: *X*) is denoted as *T* (eqs. 5-6) and the trial-by-trial covariance (here: *V*) is denoted as *U* (eq. 7). In previous work on ITEMs, we have shown that 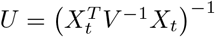 where *X* is the trial-wise design matrix and *V* is the scan-by-scan covariance (Soch et al., 2020, eq. 7).

3 The simplest way of combining these predictions is, for each trial, to return the experimental condition which is compatible with each predicted experimental factor, e.g. 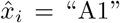 when 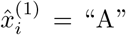 and 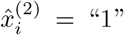 etc. A more sophisticated approach can be applied when the decoding algorithm (e.g. logistic regression) returns trial-wise probabilities such as 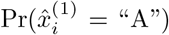 and 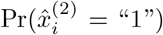. These probabilities can then be used as marginal probabilities which imply the trial-wise probability of each (joint) experimental condition, if experimental design dimensions are regarded as independent, i.e. 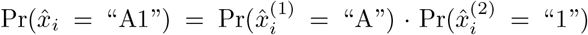. For details, see Supplementary Section 1 and Supplementary Appendix B.

4 https://www.narps.info/

5 https://openneuro.org/datasets/ds001734/

6 https://openneuro.org/datasets/ds001734/versions/1.0.5/file-display/derivatives:fmriprep:logs:CITATION.html

7 https://github.com/JoramSoch/ITEM

8 www.bccn-berlin.de/tdt

## Notes

### Competing Interest Statement

The authors have declared no competing interest.

