## Supplementary Material for "Indirect decoding of behavior from brain signals via reconstruction of experimental conditions"

**Abstract**

In this supplementary information, we share additional material about indirect response decoding (iRD) including (i) an empirical pilot study using a smaller data set conducted before the main analysis (see Section 1); (ii) supplementary results in the form of balanced accuracies and prevalence inference for the main analysis (see Section 2); (iii) a supplementary discussion on philosophical aspects of relating mind, brain and behavior (see Section 3); and (iv) a supplementary appendix detailing some of the mathematics behind iRD (see Section 4).

**5 References** **20**

### 1 Empirical pilot study

Before the data analysis reported in the paper (see Sections 3 and 4 in the main paper), we performed a pilot study using a smaller data set and searchlight-based decoding (Soch and Haynes, 2020) in order to investigate the relative ranking of sbRD, dRD and iRD.

#### 1.1 Methods

We analyzed the example data set (Hebart and Görgen, 2015) of *The Decoding Toolbox* (TDT; Görgen et al., 2012; Hebart et al., 2015). In this experiment, subjects[1] were cued to focus on a specific stimulus property (color or direction), then observed a stimulus (red or green dots moving up or down) and gave a binary behavioral response (see Figure S1A). Participants had to respond with the left or right button, depending on the presented response mapping that indicated which stimulus features required the left and right button (see Figure S1B). Thus, there are five experimental dimensions ($X$; cue, stimulus color and direction, color and direction requiring left button press) and one behavioral dimension ($Z$; left vs. right button press), each having two levels.
First, general linear models (GLMs) were specified modelling (i) cue phases, (ii) stimulus phases, (iii) response mapping phases and (iv) actual response phases. Then, trial-wise response amplitudes were estimated using the ITEM toolbox (Soch et al., 2020) for (a) the stimulus phase ($Y_1$) and (b) the response phase ($Y_2$) of all $t = 32$ trials in each of the $S = 8$ fMRI sessions, resulting in $S \times t = 256$ observations per subject[2].
*Stimulus-based response decoding* $Z = g(X)$ was performed by estimating transition probabilities from each of the $p = 2^5 = 32$ experimental conditions to each of the $q = 2$ response options (see Figure S2A). This can be done by simply regressing the indicator matrix $Z$ against the indicator matrix $X$ (see Appendix A), such that $p_{jk}$ is the probability of moving from condition $j$ to response $k$.
For *direct response decoding* $Z = h(Y_2)$, behavioral responses were directly decoded from response-related fMRI signals using logistic regression (target: left vs. right button press; features: fMRI signals during response phase). This allows to calculate the probability that a particular left-out trial results in a left or right button press. By thresholding this probability (at 0.5), a prediction can be made.
For *indirect response decoding* $Z = g\left(f^{-1}(Y_1)\right)$, experimental conditions were decoded from stimulus-related fMRI signals using logistic regression (targets: red vs. green, up vs. down; features: fMRI signals during stimulus phase). In left-out trials, predictions for stimulus color and direction were combined with actual cue and response mapping to yield a constructed design $\hat{X}$. In this combination, we made use of the fact that marginal probabilities provided by logistic regression (e.g. $\mathrm{Pr}(x_i^{(1)} = \text{“red”} | y_i)$ and $\mathrm{Pr}(x_i^{(2)} = \text{“up”} | y_i)$) can be combined into a joint probability (e.g. $\mathrm{Pr}(x_i = \left\{\text{“red”}, \text{“up”}\right\} | y_i)$) when experimental dimensions are assumed independent (see Appendix B). Finally, out-of-sample behavioral responses were predicted by submitting the out-of-sample predicted design to the psychobehavioral model estimated from within-sample data: $\hat{Z}_{\text{test}} = \hat{g}_{\text{train}}\left(\hat{X}_{\text{test}}\right)$.

[1] Two subjects performed the experiment, labeled as “S1” and “S2” in Figure S2.

[2] If a subject did not respond during a trial, no response-related activation (but condition-related activation) was estimated and this trial was removed from all subsequent behavioral decoding analyses (but not from experimental design decoding). S1 responded in all 256 trials and S2 responded in 252 trials.

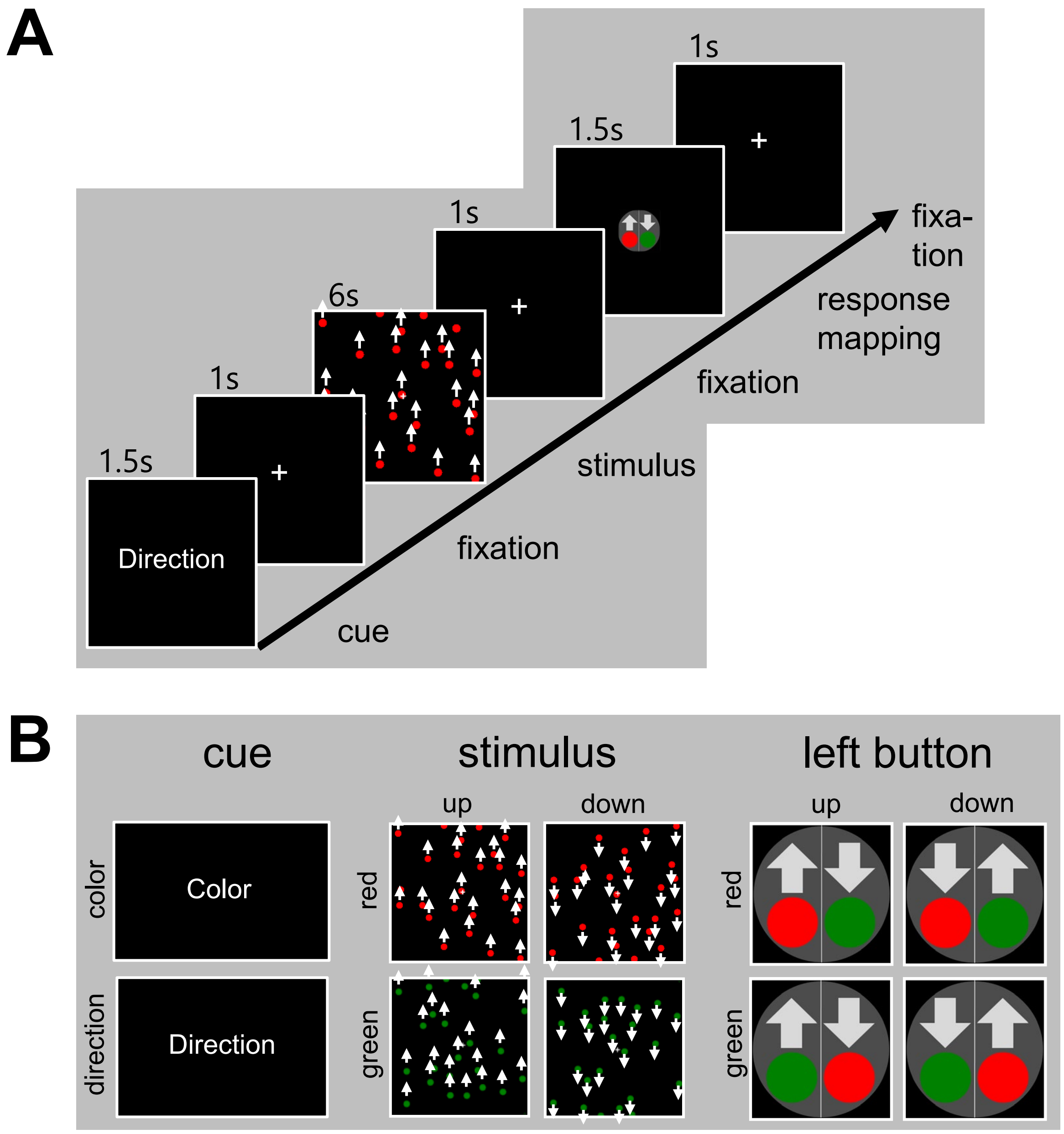

Figure S1: *Experimental design of decision-making task.* **(A)** Each trial of the experiment consisted of a cue phase (indicating which stimulus property to focus on), a stimulus phase (presenting the stimulus) and a response (mapping) phase (requiring a behavioral response), all separated by 1 s fixation phases. **(B)** The experimental design consisted of five two-level factors (cue, stimulus color and direction, color and direction requiring left button press), resulting in $2^5 = 32$ experimental conditions, each of which was presented once during each of the 8 sessions, resulting in $8 \times 32 = 256$ trials.

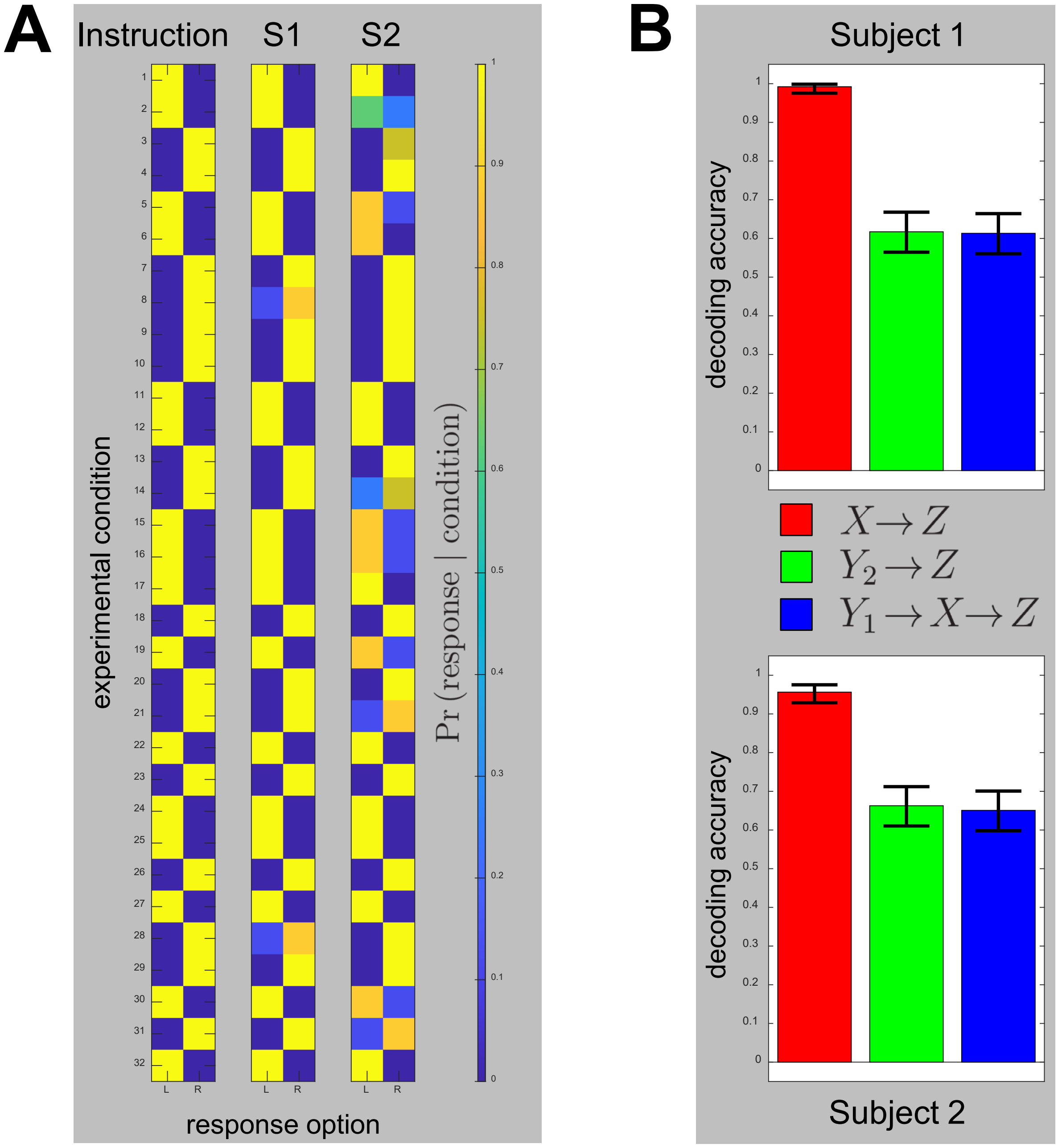

Figure S2: *Behavioral responses and decoding accuracies.* **(A)** Instructed task and behavioral data. The instruction is visualized as an indicator matrix mapping from conditions ($y$-axis) to responses ($x$-axis). Subject S1 performed almost perfectly whereas subject S2 made a few mistakes. **(B)** Decoding accuracies for both subjects. Accuracy is calculated across all sessions when predicting button presses using stimulus-based ($X \to Z$; red), direct ($Y_2 \to Z$; green) and indirect ($Y_1 \to X \to Z$; blue) response decoding. Error bars on decoding accuracies represent 90% binomial confidence intervals ($n = 256$ for S1 and $n = 252$ for S2, see fn. 2).

#### 1.2 Results

Prediction of experimental design and behavioral response variables from measured fMRI signals was performed via searchlight-based decoding using a searchlight radius of $r = 6$ mm, leading to a typical number of 71 voxels per searchlight.

To make dRD and iRD comparable, selection of the most informative searchlights was performed by maximizing within-sample decoding accuracy and reporting results as out-of-sample decoding accuracies in the selected searchlights (Soch and Haynes, 2020, Fig. 1B). For iRD, decoding accuracy was separately maximized for the decoded design variables (stimulus color and stimulus direction) and the resulting single-searchlight predictions were combined into a reconstructed experimental design (cf. Section 2.6 of the main manuscript and see Section 1.1 in this supplementary material).

When looking at the empirical results in Figure S2, we observe two main results:

- First, sbRD achieves near-perfect decoding accuracy and strongly outperforms dRD and iRD. This is due to the fact that the experiment was an instructed task and subjects behaved closely to this instruction (see Figure S2A).
- Second, we obtain statistically indistinguishable performance of dRD and iRD in both subjects (see Figure S2B). This mirrors results obtained in the main manuscript, especially when decoding discrete $Z$ from continuous $X$ (see Figure 4).

### 2 Supplementary Results

In these supplementary results, we report alternative views of the group-level data, namely (i) box plots of balanced accuracies (rather than decoding accuracies) and (ii) bar plots of prevalence estimates (rather than average estimates).

#### 2.1 Balanced accuracies

Because subjects' responses are not experimentally controlled, there can be an unequal number of responses across categories (e.g. left and right button; here: accept/reject). In such a case of class imbalance, it is advisable to also compute balanced accuracies Brodersen et al., 2010) next to decoding accuracies.

For all analyses in which discrete (design or response) variables were predicted (see Table 4 in the main manuscript), we provide box plots of balanced accuracies in Figures S3, S4 and S5 which are mirroring Figures 4, 5 and 6 from the main manuscript. For all analyses in which continuous variables were predicted, we report the same correlation coefficients as in the main figures for comparison purposes.

While balanced accuracies are generally lower than decoding accuracies for dRD as well as iRD (cf. Figure 4 vs. Figure S3), this does not seem to affect the two main results that (i) dRD and iRD both allow for above-chance behavioral decoding and (ii) dRD only mildly outperforms iRD in most analysis scenarios.

Note that the balanced accuracies for discrete experimental design variables do not differ from the corresponding decoding accuracies (cf. Figure 6 vs. Figure S5), consistent with the fact that there are no class imbalances for design variables (i.e. gain and loss) which were experimentally controlled to be evenly distributed.

#### 2.2 Prevalence inference

Following recent advances in Bayesian inference of population prevalence (Ince et al., 2020), we also computed prevalence estimates for each decoding algorithm (per each analysis strategy), i.e. estimates of the proportion of subjects showing the effect which the successful decoding is based on.

To this end, we determined single-subject $p$-values (i) for decoding accuracies (using binomial confidence intervals) in the case of discrete predictions and (ii) for correlation coefficients (using Student's $t$-test) in the case of continuous predictions. Then, the number of significant results was determined for each experimental group and submitted to the "bayesian-prevalence" package.[3]

The resulting prevlance estimates are given in Figures S6, S7 and S8 which are mirroring Figures 4, 5 and 6 from the main manuscript. Overall, these results support the notion that effects which dRD is based on are slightly more prevalent than effects which iRD is based on (see Figure S6).

[3]From the MATLAB code supplied on GitHub (https://github.com/robince/bayesian-prevalence), we used `bayesprev_map` to compute maximum-a-posteriori (MAP) estimates and `bayesprev_hpdi` to compute highest posterior density intervals (HPDI). Input parameters were set to $\alpha = 0.05$ and $\beta = 1$ as well as $p = 0.9$ (for 90% HPDIs).

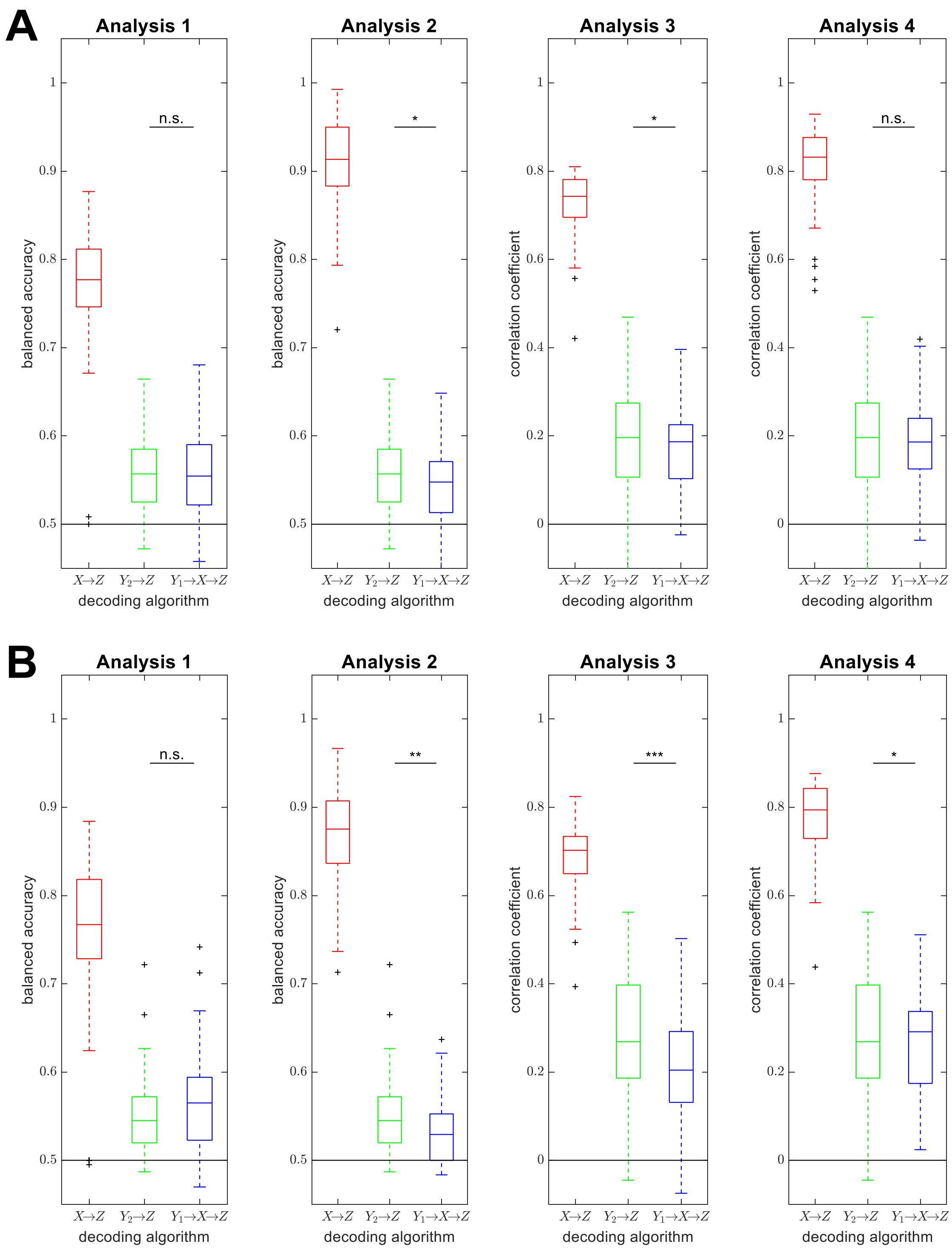

Figure S 3: *Balanced accuracies as a function of decoding method.* Performances of stimulus-based ($X \rightarrow Z$; red), direct ($Y_2 \rightarrow Z$; green) and indirect ($Y_1 \rightarrow X \rightarrow Z$; blue) response decoding are visualized using box plots for four analysis types, separately for the **(A)** equal range and the **(B)** equal indifference condition. Direct and indirect response decoding are tested against each other using a two-tailed paired t-test. Abbreviations: n.s. = not significant; * $p < 0.05$, ** $p < 0.01$, *** $p < 0.001$. (This figure mirrors Figure 4 from the main manuscript.)

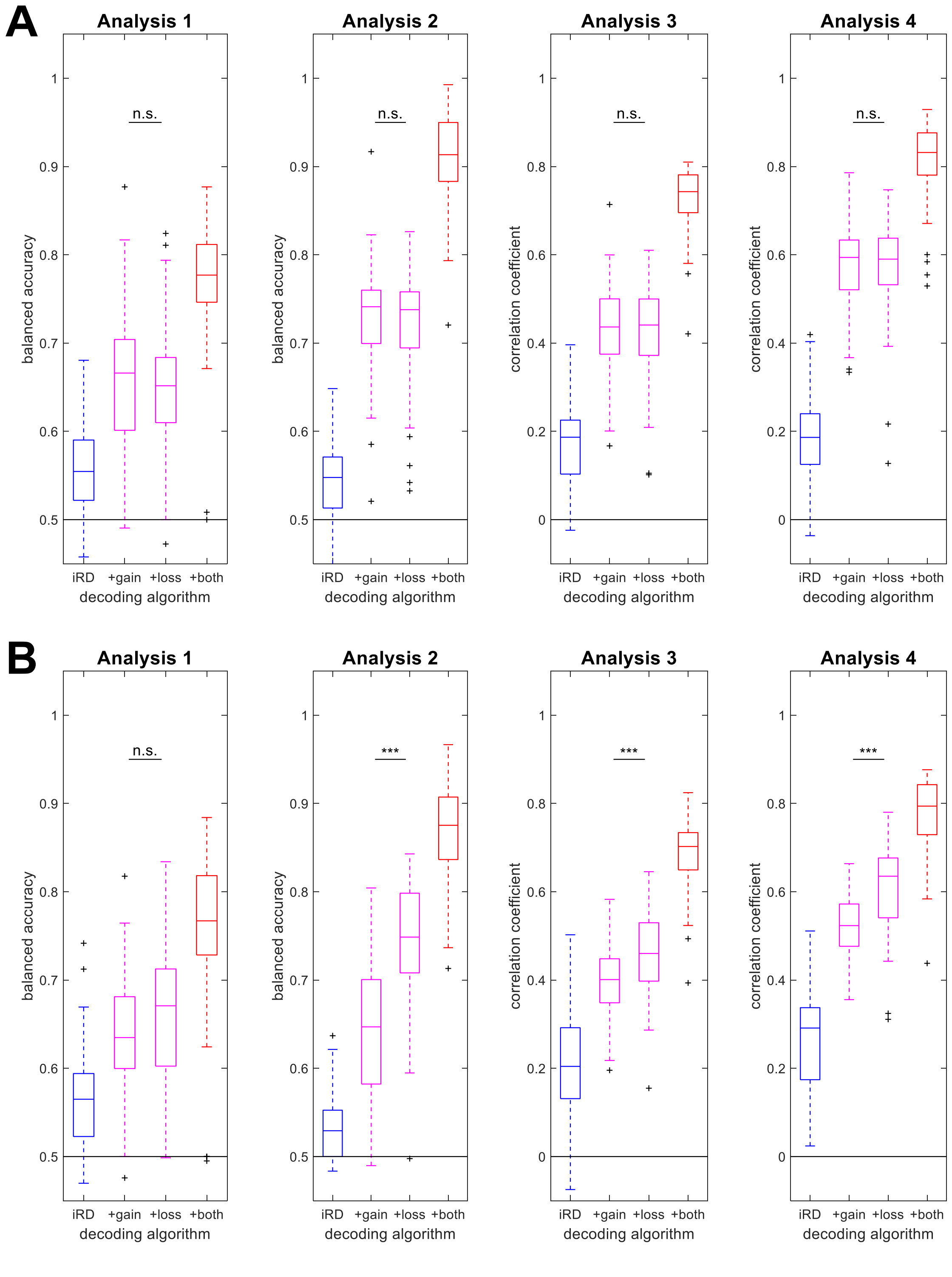

Figure S 4: *Balanced accuracies as a function of information supply.* Performances of indirect response decoding (iRD), depending on whether both experimental dimensions were reconstructed from the data (blue, left), whether one of the experimental dimensions was not reconstructed, but supplied to the behavioral model (magenta, middle) or whether both experimental dimensions were known (red, right). The layout follows the one of Figure S3 and gives results for four analysis types, separately for the **(A)** equal range and the **(B)** equal indifference condition. Box plots colored in blue and red are identical to those labeled as "$Y_1 \rightarrow X \rightarrow Z$" and "$X \rightarrow Z$" on Figure S3. (This figure mirrors Figure 5 from the main manuscript.)

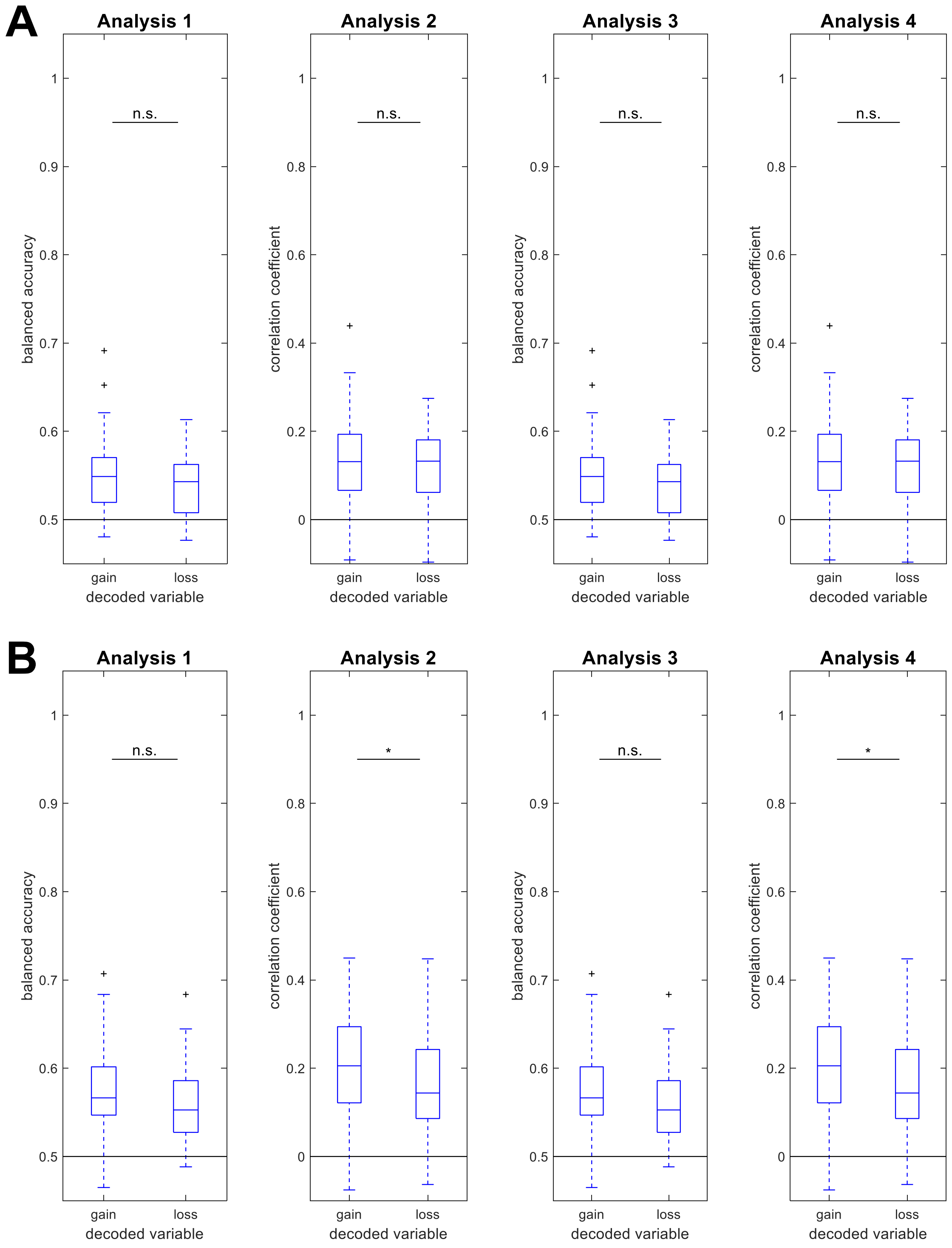

Figure S5: *Balanced accuracies for experimental design variables.* Performances of reconstructing the experimental design from measured fMRI signals. The layout follows the one of Figure S3 and gives results for four analysis types, separately for the **(A)** equal range and the **(B)** equal indifference condition. (This figure mirrors Figure 6 from the main manuscript.)

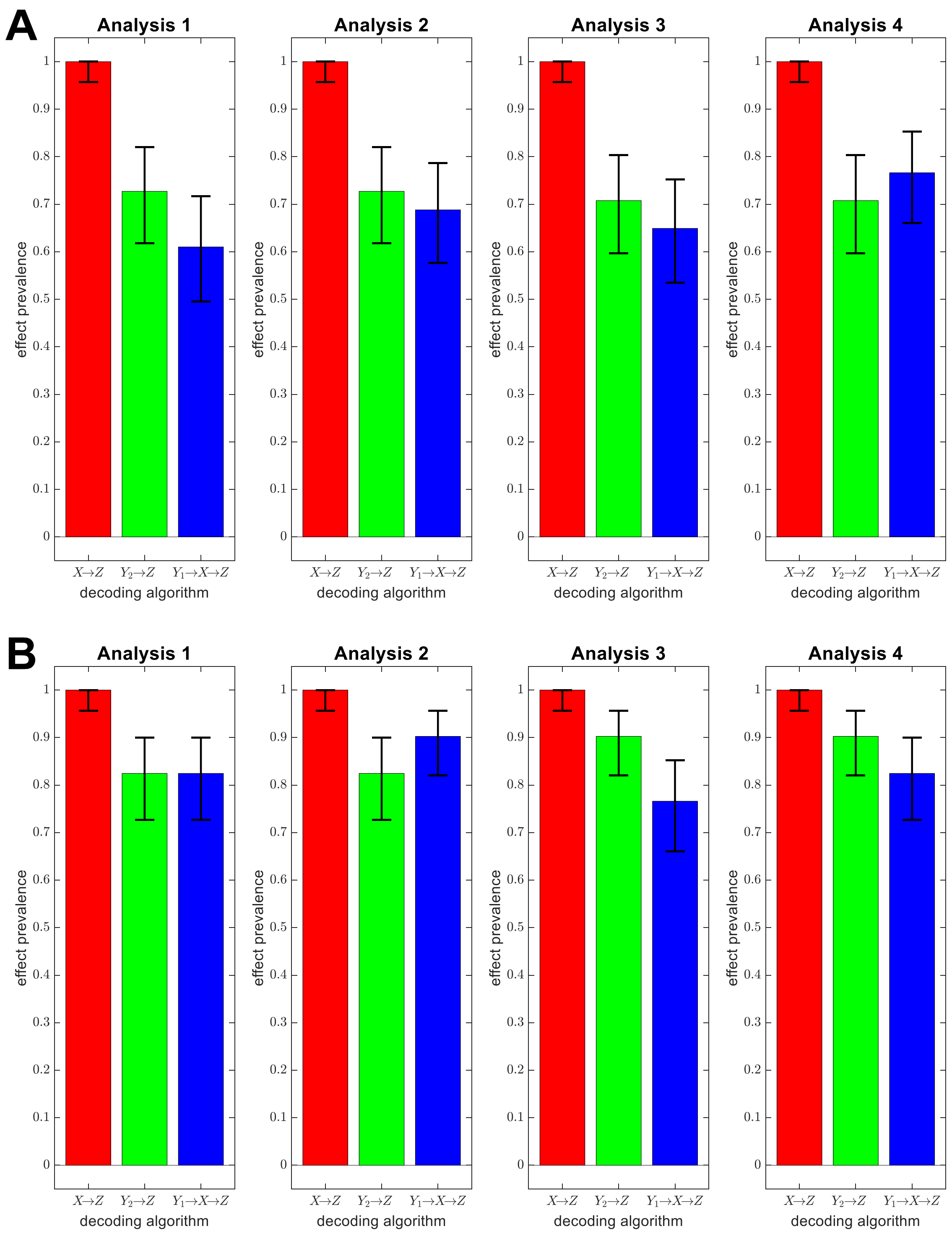

Figure S 6: *Prevalence estimates as a function of decoding method.* Effect prevalences for stimulus-based ($X \to Z$; red), direct ($Y_2 \to Z$; green) and indirect ($Y_1 \to X \to Z$; blue) response decoding are given as maximum-a-posteriori (MAP) estimates and highest posterior density intervals (HPDI) for four analysis types, separately for the **(A)** equal range and the **(B)** equal indifference condition. (This figure mirrors Figure 4 from the main manuscript.)

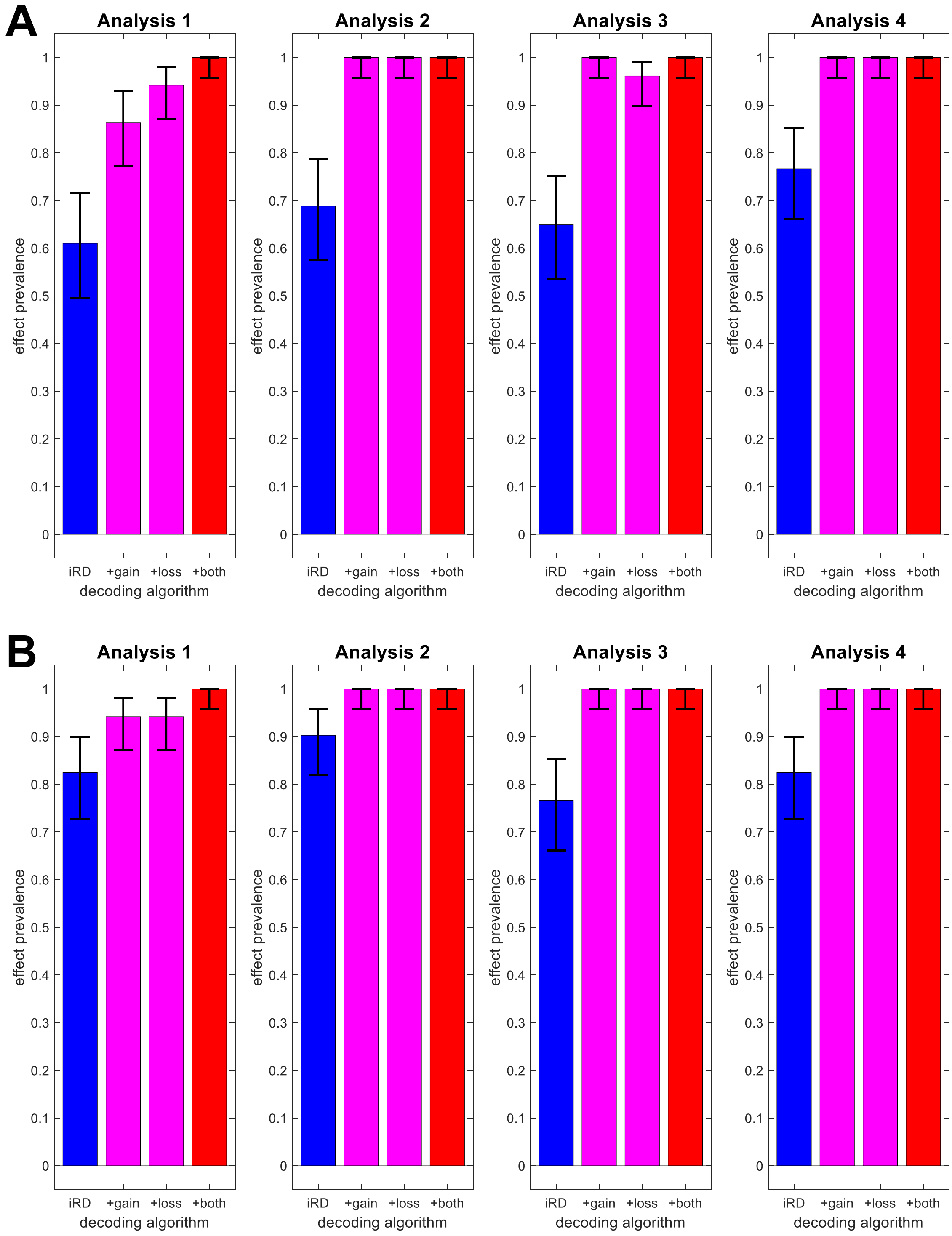

Figure S7: *Prevalence estimates as a function of information supply.* Effect prevalences for indirect response decoding (iRD), depending on whether both experimental dimensions were reconstructed from the data (blue, left), whether one of the experimental dimensions was not reconstructed, but supplied to the behavioral model (magenta, middle) or whether both experimental dimensions were known (red, right). The layout follows the one of Figure S6 and gives results for four analysis types, separately for the **(A)** equal range and the **(B)** equal indifference condition. Bar plots colored in blue and red are identical to those labeled as "$Y_1 \rightarrow X \rightarrow Z$" and "$X \rightarrow Z$" on Figure S6. (This figure mirrors Figure 5 from the main manuscript.)

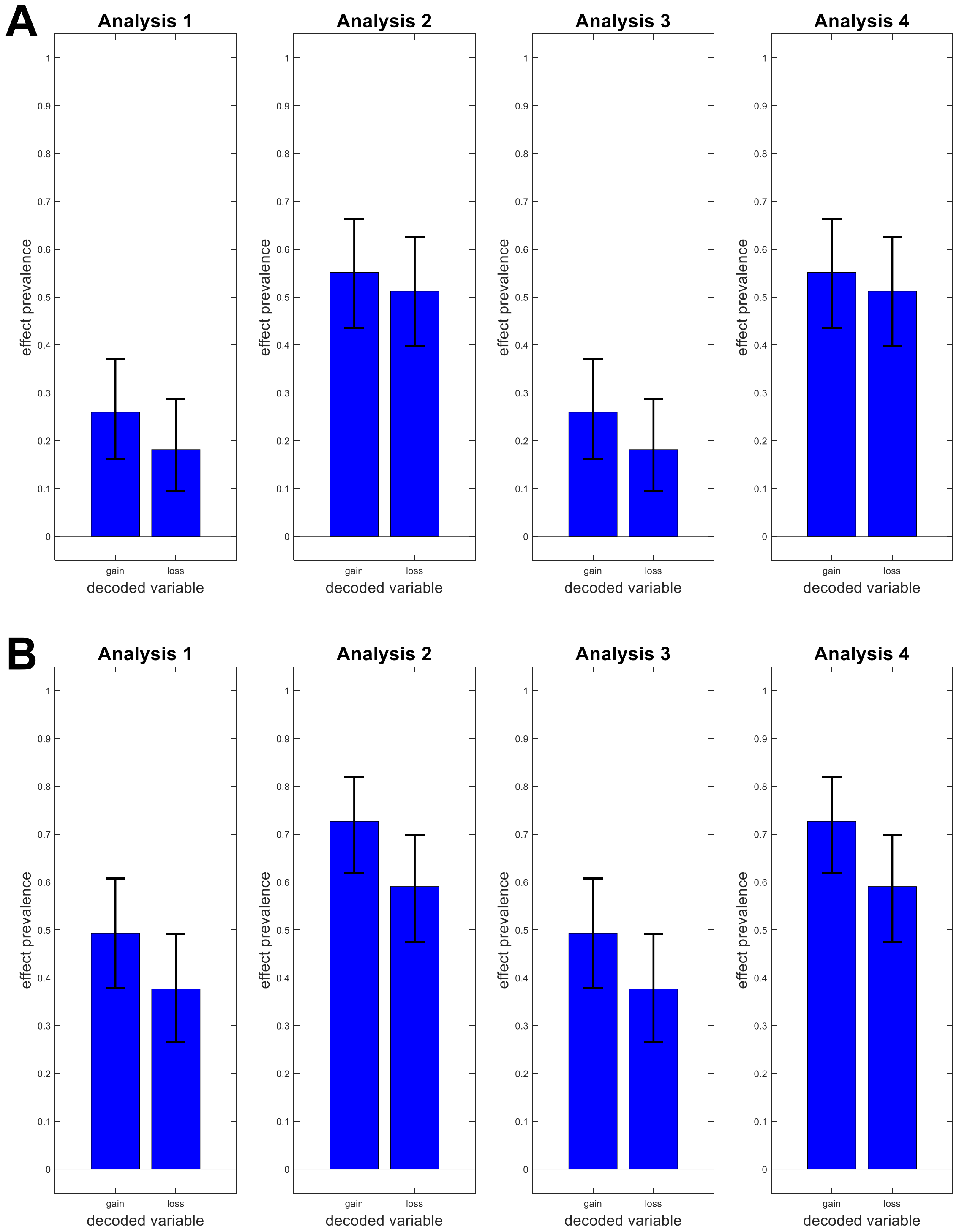

Figure S8: *Prevalence estimates for experimental design variables.* Effect prevalances for reconstructing the experimental design from measured fMRI signals. The layout follows the one of Figure S6 and gives results for four analysis types, separately for the **(A)** equal range and the **(B)** equal indifference condition. (This figure mirrors Figure 6 from the main manuscript.)

### 3 Supplementary Discussion

In this supplementary discussion, we elaborate on some philosophical aspects of indirect response decoding, especially competing models of mind, brain and behavior that might be presupposed by this technique (see Figure S9).

#### 3.1 Extensional identity of sensory input and cognitive states

First, we want to raise awareness for the fact that, in neuroimaging experiments and data analysis, cognitive states are fully operationalized via the sensory input or experimental stimuli that are assumed to elicit them in human subjects.

For example, if we *stimulate* subjects with red moving circles versus blue stationary squares, they are assumed to have *perceptions* of color (red vs. blue), movement (moving vs. stationary) and shape (circles vs. squares). Or, if specific amounts of money to gain or lose are *displayed* on the screen, subjects are assumed to perform *reward processing* operations reflecting these gains and losses (cf. Figure 3).

We refer to this as "extensional identity of experimental stimulation (or sensory input) and cognitive states" (denoted by the symbol "$\triangleq$" ("conforms to") in Figure S9A). Here, *extensional identity* implies an identity of *classes*, whereas *intensional identity* would imply an identity of *concepts*.[4]

In other words, while the words "display of red dots" and "perception of red color" of course *mean* different things – the latter is a mental state of a human subject, whereas the former is a physical state of a presentation device –, they *refer to* exactly the same set of situations in the context of a neuroimaging experiment. Similarly, "being offered to win 10 units" and "processing a potential reward of 10 units" represent different *concepts* – one consisting in a social or linguistic act, the other consisting in a mental operation –, but constitute the exact same *class* of events in the experimental design underlying the NARPS data set (see Secton 3.1).

In conclusion, this means that neuroimaging data analysis makes no practical difference between experimental stimulation (or sensory input) and the cognitive states it elicits. Decoding actually presented stimulus color is (extensionally) equivalent to decoding a subject's perception of color. And reconstruction of actually applied gain and loss values is (extensionally) equivalent to reconstruction of a subject's reward expectations.

#### 3.2 Neural signals as an epiphenomenon to cognitive states?

When attempting to isolate the system of variables underlying indirect response decoding, a superficial treatment could lead to the following conceptualization: *Sensory input* leads to *cognitive states* $X$ (and is extensionally identical with them – see previous section) and cognitive states themselves translate into *behavioral responses* $Z$. *Neural signals* $Y$ are a side-effect of cognitive states (see Figure S9A), but allow for a mapping to those cognitive

[4] Simplified, extensional identity of A and B can be understood as full coincidence (or sufficient correlation) of A-objects and B-objects and intensional identity can be understood as synonymity of the symbols "A" and "B" (cf. Ros, 2005, pp. 227-232). Intensional identity implies extensional identity (e.g. "bachelors" and "unmarried, non-divorced, non-widowed adult men"), but extensional identity does not imply intentional identity (e.g. "living beings with a heart" and "living beings with a kidney").

states (inverse neurophysiological model, $f^{-1}$) and behavioral responses (conventional response decoding, $h$).
Therefore, neural signals appear to be no more than an *epiphenomenon to cognitive states*, with the relevant causal chain completely established by sensory input, cognitive states and behavioral output via, for example, a stimulus-response mapping (psychobehavioral model, $g$). This is a paradoxical conclusion, given that, if at all, mental (cognitive) states are usually thought to be an epiphenomenon of physical (neural) states in philosophy of mind, not vice versa (Beckermann, 2008, ch. 3.1.1).
However, we argue that this impression is more or less an artifact of the indirect response decoding (iRD) approach. Because cognitive states $X$ are the target for prediction from neural signals $Y$ (iRD's 1st step) and then form the source for prediction of behavioral responses $Z$ (iRD's 2nd step), they appear in the center of the framework by construction (but other frameworks are possible – see next section).

#### 3.3 Epistemological pluralism with respect to cognitive states

Specifically, we want to suggest an epistemologically consistent framework in which both, neural signals $Y$ and behavioral responses $Z$, are in some sense epiphenomena to cognitive states $X$. This conception assumes a continuous stream of cognitive states which is influenced by prior knowledge (e.g. task instructions) as well as sensory input (i.e. experimental stimulation) and itself influences our measurements of neural and behavioral outputs (see Figure S9B).
We refer to this conception as "epistemological pluralism with respect to cognitive states", because measured brain signals and acquired behavioral data would be regarded as different, but equal levels of description of the same underlying phenomena (i.e. cognitive states). Note that these descriptions are not restricted to neural and behavioral observables, but in theory could also be based on e.g. physiological (e.g. respiratory signals) or hormonal (e.g. neuropeptide expression) observations.
In indirect response decoding, both sets of observed data, $Y$ and $Z$, are thought to emerge from $X$ by generative models mapping from cognitive states to neural signals (neurophysiological model, $f$) or behavioral responses (psychobehavioral model, $g$). Additionally, the two levels of description can be connected with each other by a function expressing $Z$ in terms of $Y$ (conventional response decoding, $h$).
These three functions can also be embedded into another, less epistemological framework which regards the brain as an input-state-output system, where (i) input into the brain is considered to be controlled via experimental stimulation $X$; and (ii) internal states and system outputs are indirectly accessible via acquired neural signals $Y$ and recorded behavioral responses $Z$ (see Figure S9C).
This approach tracks the complete causal chain from $X$ over $Y$ to $Z$ using one mapping $f$ from controlled experimental stimulation (extensionally identical to cognitive states – see Section 3.1) to measured neural signals and another mapping $h$ from measured brain data to observed behavioral responses. The third mapping $g$ would then be regarded as a way to circumvent variables of type $Y$.[5]

[5] In future work, we want to employ this framework to place variables $X$, $Y$, $Z$ to their respective positions in the assumed causal chain of cognitive processing which would allow understanding the role of brain regions and networks as a mediator between sensory input and behavioral output.

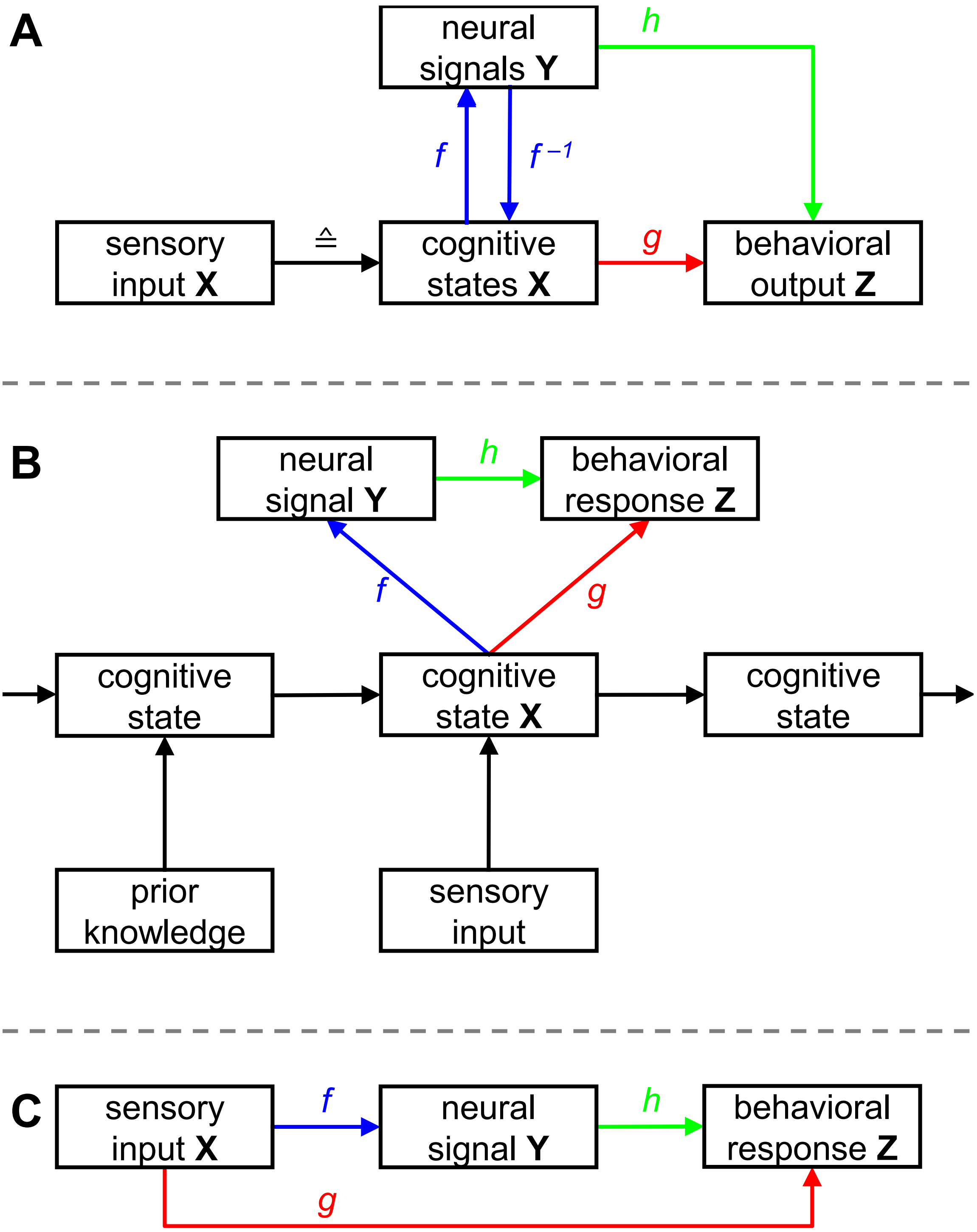

Figure S9: *Competing models of mind, brain and behavior.* The variables $X$, $Y$, $Z$ and the functions $f$, $g$, $h$ are similar to those in Figure 1 of the main manuscript. **(A)** Superficial ontology underlying indirect response decoding. Sensory input causes cognitive states causing behavioral output with neural signals as an epiphenomenon of cognitive states. **(B)** Alternative interpretation of indirect response decoding. A stream of cognitive states is influenced by prior knowledge and sensory input and manifests itself in neural signals and behavioral responses. **(C)** Envisaged ontology for future work. Sensory input causes neural signals causing behavioral responses, representing the commonly assumed causal chain of cognitive processing.

### 4 Supplementary Appendix

#### A Indicator matrices and transition probabilities

Suppose that the experimental design is given as an $n \times p$ indicator matrix $X$ ($j = 1, \ldots, p$ experimental conditions) and that the behavioral responses are given as an $n \times q$ indicator matrix $Z$ ($k = 1, \ldots, q$ response options), for a number of observations ($i = 1, \ldots, n$). Further, let the true and estimated transition probabilities be defined as:

$$\begin{aligned} p_{jk} &= \mathrm{Pr}(z_k = 1 | x_j = 1) \\ \hat{p}_{jk} &= \frac{\sum_{i=1}^{n} z_{ik} \cdot x_{ij}}{\sum_{i=1}^{n} x_{ij}} \; . \end{aligned} \tag{A.1}$$

Then, we can show that the following holds:

**Theorem 1:** Regressing $Z$ against $X$ using ordinary least squares (OLS) results in empirical transition probabilities:

$$\hat{P} = (X^\mathrm{T} X)^{-1} X^\mathrm{T} Z \; . \tag{A.2}$$

**Proof:** $X$ and $Z$ are indicator matrices, so we have $x_{ij} \in \{0, 1\}$ and $z_{ik} \in \{0, 1\}$ for all $i, j, k$. Consider the OLS solution

$$(X^\mathrm{T} X)^{-1} X^\mathrm{T} Z \; . \tag{A.3}$$

Because experimental conditions are mutually exclusive[6], $X^\mathrm{T} X$ is a $p \times p$ diagonal matrix with numbers of trials in each condition on the diagonal:

$$X^\mathrm{T} X = \mathrm{diag}\left( [n_1, \ldots, n_p] \right) \; . \tag{A.4}$$

The inverse of a diagonal matrix is a diagonal matrix in which each entry on the diagonal has been inverted:

$$(X^\mathrm{T} X)^{-1} = \mathrm{diag}\left( [1/n_1, \ldots, 1/n_p] \right) \; . \tag{A.5}$$

Because $X$ and $Z$ are indicator matrices, $X^\mathrm{T} Z$ is a $p \times q$ matrix the $(j, k)$-th entry of which is given by the number of trials in which condition $j$ was active and response $k$ was given:

$$(X^\mathrm{T} Z)_{jk} = \sum_{i=1}^{n} z_{ik} \cdot x_{ij} \; . \tag{A.6}$$

Following the rules of matrix multiplication, the product of $(X^\mathrm{T} X)^{-1}$ and $X^\mathrm{T} Z$ is a $p \times q$ matrix, the $(j, k)$-th entry of which is given by:

$$\left[ (X^\mathrm{T} X)^{-1} X^\mathrm{T} Z \right]_{jk} = \frac{1}{n_j} \cdot \sum_{i=1}^{n} z_{ik} \cdot x_{ij} = \frac{\sum_{i=1}^{n} z_{ik} \cdot x_{ij}}{\sum_{i=1}^{n} x_{ij}} = \hat{p}_{jk} \; . \tag{A.7}$$

[6] See: https://statproofbook.github.io/D/exc.

#### B Logistic regression and conditional probabilities

Suppose that we have estimated logistic regression models from training data which supply us with conditional probabilities for trial types in test data

$$\begin{aligned} &\mathrm{Pr}(x_i^{(l)} = 1|y_i), \;\; l = 1, \ldots, d \\ &\mathrm{Pr}(x_i^{(l)} = 0|y_i) \;\; = 1 - \mathrm{Pr}(\hat{x}_i^{(l)} = 1|y_i) \end{aligned} \tag{B.1}$$

where $i$ is the index of a left-out trial, $l$ is the index of a (binary) experimental dimension (e.g. red vs. green or up vs. down) and $d$ is the number of factors.
Further, assume that we have fitted a psychobehavioral model mapping from conditions to resposes using transition probabilities (see Appendix A)

$$\hat{p}_{jk} = \mathrm{Pr}(z_{ik} = 1|x_{ij} = 1) \tag{B.2}$$

where $j$ and $k$ index experimental condition and response option, respectively.
Then, we can show that the following holds:

**Theorem 2:** The estimated conditional probabilities of observing the response $k$ in trial $i$, given the measured signals in this trial, are

$$\mathrm{Pr}(z_{ik} = 1|y_i) = \hat{x}_i \, \hat{p}_{\bullet k} \tag{B.3}$$

where $\hat{x}_i$ is a $1 \times p$ vector of joint (condition) probabilities $\mathrm{Pr}(x_{ij} = 1|y_i)$ in trial $i$, obtained from marginal (dimension) probabilities $\mathrm{Pr}(x_i^{(l)} = 1|y_i)$ in this trial [eq. B.1], and $\hat{p}_{\bullet k}$ is the $k$-th column of the estimated transition probability matrix $\hat{P}$ [eq. B.2].

**Proof:** In a binary factorial design, each experimental condition $j = 1, \ldots, p$ is characterized by a $1 \times d$ indicator vector $f_j$, signifying whether each factor is "on" or "off" for this condition in a particular trial:

$$x_{ij} = 1 \quad \Leftrightarrow \quad x_i^{(1)} = f_{j1} \;\wedge\; \ldots \;\wedge\; x_i^{(d)} = f_{jd} \; . \tag{B.4}$$

If experimental dimensions can be considered statistically independent – which can be assumed for a controlled experimental design with balanced trial numbers –, their joint probability is the product of the marginal probabilities[7]:

$$\begin{aligned} \hat{x}_{ij} &= \mathrm{Pr}(x_i^{(1)} = f_{j1} \;\wedge\; \ldots \;\wedge\; x_i^{(d)} = f_{jd}|y_i) = \prod_{l=1}^{d} \mathrm{Pr}(x_i^{(l)} = f_{jl}|y_i) \\ &= \mathrm{Pr}(x_{ij} = 1|y_i) = \prod_{l=1}^{d} \mathrm{Pr}(x_i^{(l)} = f_{jl}|y_i) \; . \end{aligned} \tag{B.5}$$

[7]See: https://statproofbook.github.io/D/ind.

Using the law of marginal probability[8] and the law of conditional probability[9], the probability of response $k$ in trial $i$, given the measured signals in this trial, can be written as

$$\Pr(z_{ik} = 1|y_i) = \sum_{j=1}^{p} \Pr(z_{ik} = 1|x_{ij} = 1) \Pr(x_{ij} = 1|y_i) \tag{B.6}$$

which, using the results in eqs. B.2 and B.5 can be written as

$$\begin{aligned} \Pr(z_{ik} = 1|y_i) &= \sum_{j=1}^{p} \hat{p}_{jk} \, \hat{x}_{ij} \\ &= \sum_{j=1}^{p} \hat{x}_{ij} \, \hat{p}_{jk} \\ &= \begin{bmatrix} \hat{x}_{i1} & \dots & \hat{x}_{ip} \end{bmatrix} \cdot \begin{bmatrix} \hat{p}_{1k} \\ \vdots \\ \hat{p}_{pk} \end{bmatrix} \\ &= \hat{x}_i \, \hat{p}_{\bullet k} \; . \end{aligned} \tag{B.7}$$

[8]See: https://statproofbook.github.io/D/prob-marg.
[9]See: https://statproofbook.github.io/D/prob-cond.

### C Indirect response decoding using multivariate GLMs

In this section, we want to give a mathematical suggestion why iRD achieved comparable performance with dRD in our data analyses. For simplicity, we will assume just one set of fMRI signals $Y$ (rather that stimulus-related vs. response-related signals).
Suppose that the neurophysiological model (NPM) $f$ mapping from experimental design $X$ to measured signals $Y$ is a multivariate general linear model (GLM)

$$Y = XB + E, \; E \sim \mathcal{MN}(0, V, \Sigma_y) \tag{C.1}$$

and that the psychobehavioral model (PBM) $g$ mapping from experimental design $X$ to behavioral responses $Z$ is also a multivariate GLM

$$Z = XP + N, \; N \sim \mathcal{MN}(0, I_t, \Sigma_z) \; . \tag{C.2}$$

For example, this would be the case, if we used transformed encoding models (Soch et al., 2020, eq. 10) as neurophysiological models and multivariate linear regression as psychobehavioral models. It would also be the case, if we used voxel-wise univariate GLMs for the mapping $X \rightarrow Y$ and simple transition probabilities for the mapping $X \rightarrow Z$, as in the main paper (cf. Analysis 1 in Section 3.4).
Our goal will now be to represent $Z$ as a function of $Y$ using $X$ – as in the indirect response decoding framework (see Section 2) – and, on this basis, suggest an explanation for the fact that iRD achieves equal decoding performance with dRD, despite a more complex reconstruction approach. Specifically, we will show the following:

**Theorem 3:** Given the NPM in (C.1) and the PBM in (C.2), $Z$ can be written as a multivariate GLM function of $Y$

$$Z = YM + \tilde{N}, \; \mathrm{vec}(\tilde{N}) \sim \mathcal{N}(0, \Sigma_z \otimes I_t + M^\mathrm{T} \Sigma_z M \otimes V) \tag{C.3}$$

where $M$ is a function of $B$ and $P$ and $\tilde{N}$ are normally distributed errors.

**Proof:** Given the multivariate GLM-based forward model

$$Y = XB + E, \; E \sim \mathcal{MN}(0, V, \Sigma_y) \; , \tag{C.4}$$

we know that there exists a backward model[10] (Soch et al., 2020, Theorem 4)

$$X = YW + H, \; H \sim \mathcal{MN}(0, V, W^\mathrm{T} \Sigma_y W) \tag{C.5}$$

where $W$ is a $v \times p$ matrix, such that $BW = I_p$. Such a matrix exists, if the rows of $B$ are linearly independent, in which case the optimal solution is $W = \Sigma_y^{-1} B^\mathrm{T} (B \Sigma_y^{-1} B^\mathrm{T})^{-1}$.
Now, we insert the formula for $X$ from (C.5) into the psychobehavioral model in (C.2) which yields:

$$\begin{aligned} &Z = YWP + HP + N, \\ &H \sim \mathcal{MN}(0, V, W^\mathrm{T} \Sigma_y W), \\ &N \sim \mathcal{MN}(0, I_t, \Sigma_z) \; . \end{aligned} \tag{C.6}$$

[10] See: https://statproofbook.github.io/P/iglm-dist.

With the linear transformation theorem for the matrix-normal (or multivariate normal) distribution[11], this gives:

$$HP \sim \mathcal{MN}(0, V, P^\mathrm{T} W^\mathrm{T} \Sigma_y W P) \; . \tag{C.7}$$

Substituting $WP = M$ and $HP = \tilde{H}$, we have:

$$\begin{aligned} Z &= YM + \tilde{H} + N, \\ \tilde{H} &\sim \mathcal{MN}(0, V, M^\mathrm{T} \Sigma_y M), \\ N &\sim \mathcal{MN}(0, I_t, \Sigma_z) \; . \end{aligned} \tag{C.8}$$

With the relationship between matrix-normal distribution and multivariate normal distribution[12], this gives:

$$\begin{aligned} \mathrm{vec}(\tilde{H}) &\sim \mathcal{N}(0, M^\mathrm{T} \Sigma_y M \otimes V), \\ \mathrm{vec}(N) &\sim \mathcal{N}(0, \Sigma_z \otimes I_t) \; . \end{aligned} \tag{C.9}$$

Finally, substituting $\tilde{N} = \tilde{H} + N$, we have:

$$Z = YM + \tilde{N}, \; \mathrm{vec}(\tilde{N}) \sim \mathcal{N}(0, \Sigma_z \otimes I_t + M^\mathrm{T} \Sigma_y M \otimes V) \; . \tag{C.10}$$

**Remark:** Given that trial-wise response amplitudes from fMRI signals can be regarded as independent across trials (e.g. because inter-stimulus-intervals are high enough), i.e. $V = I_t$, the model for $Z$ further simplifies to

$$\begin{aligned} Z = YM + \tilde{N}, \; \mathrm{vec}(\tilde{N}) &\sim \mathcal{N}(0, [\Sigma_z + M^\mathrm{T} \Sigma_y M] \otimes I_t) \\ \tilde{N} &\sim \mathcal{MN}(0, I_t, [\Sigma_z + M^\mathrm{T} \Sigma_y M]) \; . \end{aligned} \tag{C.11}$$

In other words, given that the generative model assumptions in Theorem 3 are true, then $Z$ is optimally represented as a projection from $Y$ with the corresponding noise matrix $\tilde{N}$ sampled from a matrix-normal distribution with covariance components $\Sigma_z$ and $M^\mathrm{T} \Sigma_y M$. We suggest that, while dRD blindly tries to find this projection without being able to account for the covariance components, iRD is aware of those covariance components by separate estimation and sequential application of the inverse NPM in (C.5) and the forward PBM in (C.2).
Note however that this is based on the assumption that stimulus-related signals $Y_1$ and response-related signals $Y_2$ are identical – an assumption that was violated by the way data analysis was performed in the main paper (see Section 3.4) as well as supplementary material (see Section 1.1)

[11]See: https://statproofbook.github.io/P/matn-ltt.
[12]See: https://statproofbook.github.io/P/matn-mvn.
